# Adaptor protein complex 2 (AP2) participates in biogenesis and homeostasis of myelin sheaths in the central nervous system

**DOI:** 10.1101/2025.09.02.673641

**Authors:** Sophie B. Siems, Ramona B. Jung, Olaf Jahn, Martin Meschkat, Susann Michanski, Nikola Łukasik, Sophie Hümmert, Andrew O. Sasmita, Wiebke Möbius, Fritz Benseler, Nils Brose, Eva-Maria Krämer-Albers, Volker Haucke, Klaus-Armin Nave, Hauke B. Werner

## Abstract

Myelination of CNS axons requires oligodendrocytes to undergo extensive morphological changes by producing large amounts of myelin membrane with defined protein composition and structure. The formation of myelin sheaths thus involves efficient trafficking and sorting of future myelin constituents via vesicles that fuse with prospective myelin membranes by exocytotic mechanisms. However, the functional relevance of other trafficking steps in oligodendocytes for myelin biogenesis is largely unknown. Here, we followed the hypothesis that developmental myelination involves endocytic mechanisms. In this model, Golgi-derived vesicles fuse with the oligodendroglial plasma membrane, from which myelin constituents are retrieved by endocytosis into endosomal/lysosomal organelles before their final integration into the growing sheath. Considering that adaptor protein complex-2 subunit-µ (AP2M) facilitates AP2-dependent endocytosis, we recombined the *Ap2m*-gene in myelin-forming oligodendrocytes, causing both hypomyelination and specific changes in the myelin proteome. Most strikingly, lysosomal membrane proteins accumulate in the abaxonal (outermost) myelin layer, identifying this membrane as an active site for retrieving constituents from myelin sheaths. These data demonstrate that the AP2 complex serves a critical function in developmental myelination *in vivo*. Unexpectedly, we also observed pathological myelin outfoldings indicative of focal hypermyelination. Consistent with the hypothesis that this phenotype reflects impaired maintenance rather than biogenesis of myelin sheaths, recombination of the *Ap2m*-gene in oligodendrocytes of adult mice caused late-onset progressive focal hypermyelination. These results indicate that, in addition to astrocytic and microglial phagocytosis, oligodendrocytes cell-autonomously contribute to maintaining the structure of healthy myelin sheaths via AP2-dependent mechanisms.

## Introduction

Oligodendrocytes are crucial for the functions of the vertebrate central nervous system (CNS) as they provide axons with myelin sheaths that enable saltatory nerve conduction, metabolic support, ion homeostasis, and antioxidant defense (Hartline and Colman, 2007; Krämer-Albers and Werner, 2023). An individual’s normal capabilities require healthy myelin sheaths with accurate ultrastructure, as evidenced by the functional decline observed in myelin-related disorders and respective animal models (Stadelmann et al., 2019; Wolf et al., 2021; Nowacki et al., 2022). The current model of developmental myelination visualizes growing myelin sheaths as fan-shaped lamellipodia-like extensions of the oligodendroglial plasma membrane, which wrap around the axon (Snaidero et al., 2014; Edgar et al., 2021). The leading edges grow both forward (radially around the axon) and laterally (longitudinally along the axon) underneath previously generated myelin layers. Newly synthesized myelin constituents are integrated into the expanding sheath predominantly at the leading edge (Snaidero et al., 2014), while compaction of myelin membranes progresses from the abaxonal (i.e., outermost) layer towards the inner tongue with a delay of 2-3 wraps behind the leading edge (Hildebrand et al., 1993). Upon maturation, the leading edge eventually forms the adaxonal (i.e., innermost) myelin layer, which remains largely uncompacted. Considering the morphological and molecular specializations of adaxonal and abaxonal myelin membranes, myelin sheaths are highly polarized. In fact, the adaxonal myelin compartment is in cytoplasmic continuity with the oligodendroglial cell body through the otherwise compacted sheath via cytoplasmic channels (Edgar et al., 2021), including paranodal myelin and the abaxonal myelin layer. The non-compacted compartments of myelin are thought to enable the trafficking of organelles, vesicles, and molecules, not only during myelin biogenesis, but also for the maintenance of mature sheaths and their cooperation with the axons they myelinate (Chapple et al., 2024).

Myelin biogenesis involves the spatiotemporally coordinated synthesis of specialized structural myelin proteins and lipids and their transport towards the growing sheath. Indeed, the delivery of myelin constituents involves multiple mechanisms. Some transcripts are localized to myelin (Thakurela et al., 2016; Yergert et al., 2021) for local synthesis of the respective myelin protein at free ribosomes; this includes myelin basic protein (MBP) (Ainger et al., 1993; Müller et al., 2013), the second-most abundant myelin protein (Jahn et al., 2020), which mediates compaction of intracellular myelin membrane surfaces (Snaidero et al., 2014). However, most myelin membrane constituents are synthesized in the oligodendrocyte soma at the rough endoplasmic reticulum and thus require transport and sorting into oligodendroglial processes before their integration into myelin membranes. Some myelin proteins coalesce with lipids as multimolecular complexes in the endoplasmic reticulum or Golgi apparatus to be transported via vesicles into the future myelin sheath (Krämer et al., 1997) as exemplified by cholesterol and the transmembrane-tetraspan proteolipid protein (PLP) (Simons et al., 2000; Werner et al., 2013), the most abundant myelin protein (Jahn et al., 2020). Consequently, the developmental expansion of myelin membranes involves exocytotic mechanisms that mediate the fusion of endosomal/lysosomal organelles with the plasma membrane (Trajkovic et al., 2006; Feldmann et al., 2011; Lam et al., 2022).

We followed the hypothesis that myelin biogenesis may also involve endocytotic recycling. According to this model, Golgi-derived vesicles fuse with the oligodendroglial plasma membrane; future myelin constituents are then retrieved via endocytosis from the plasma membrane into endosomal/lysosomal organelles before ultimately being integrated into the growing sheath. Notably, previous work using cultured oligodendrocytes has demonstrated that some myelin proteins undergo endocytosis at least *in vitro*, including PLP, myelin-associated glycoprotein (MAG), and myelin-oligodendrocyte glycoprotein (MOG) (Trajkovic et al., 2006; Winterstein et al., 2008; Baron et al., 2015). However, it remained unknown if endocytic mechanisms indeed facilitate developmental myelination *in vivo*, and, if so, whether they affect the entire myelin membrane or only particular myelin proteins.

Here we bred and analyzed mice in which the *Ap2m*-gene that encodes the medium chain subunit of adaptor protein complex 2 (AP2µ) is recombined specifically in myelinating oligodendrocytes. The AP2 complex facilitates endocytic retrieval from the plasma membrane, including clathrin-mediated endocytosis (CME) (Mettlen et al., 2018; Azarnia Tehran et al., 2019; Chen and Schmid, 2020; López-Hernández et al., 2020; Smith and Smith, 2022). Confirming our hypothesis, our data show that the AP2 complex is required for normal developmental biogenesis and molecular composition of myelin. However, it was unexpected to find that a considerable percentage of axon/myelin profiles in *Ap2m*-mutant mice displays pathological myelin outfoldings. Hypothesizing that outfoldings reflect impaired maintenance of myelin sheaths, we assessed the consequences of tamoxifen-induced recombination of the *Ap2m*-gene in myelinating oligodendrocytes of adult mice. These mice indeed displayed normal developmental myelination, but, strikingly, late-onset progressive emergence of myelin pathology. It has been known that pathological myelin sheaths can be cleared extrinsically by phagocytosing microglia (Safaiyan et al., 2016; Pluvinage et al., 2019; Djannatian et al., 2023; Olveda et al., 2024), astrocytes (Morizawa et al., 2017; Ponath et al., 2017; Wan et al., 2022a; Yang et al., 2023) and endothelial cells (Zhou et al., 2019). Importantly, thus, our results indicate that mature oligodendrocytes contribute to resolving myelin pathology cell-autonomously, involving AP2-dependent mechanisms.

## Results

### Oligodendrocytes and myelin comprise constituents of the AP2 complex

We followed the hypothesis that adaptor protein complex 2 (AP2) participates in myelination of the CNS. To pursue this hypothesis we first determined if oligodendrocytes express mRNAs encoding the subunits of the heterotetrameric AP2 complex and of the clathrin coat. Indeed, according to a previously published RNA-Seq database of cell types sorted from the cortices of mice (GEO dataset GSE52564 (Zhang et al., 2014)), *Ap2a1*, *Ap2a2*, *Ap2b1*, *Ap2m1*, *Ap2s1*, *Clta*, *Cltb*, and *Cltc* transcripts were detected in oligodendrocyte precursor cells (OPC, containing 5% of microglial contamination according to Zhang et al., 2014), newly formed oligodendrocytes (NFO), and myelinating oligodendrocytes (MOL) (**Fig. 1a**).

**Fig. 1.**
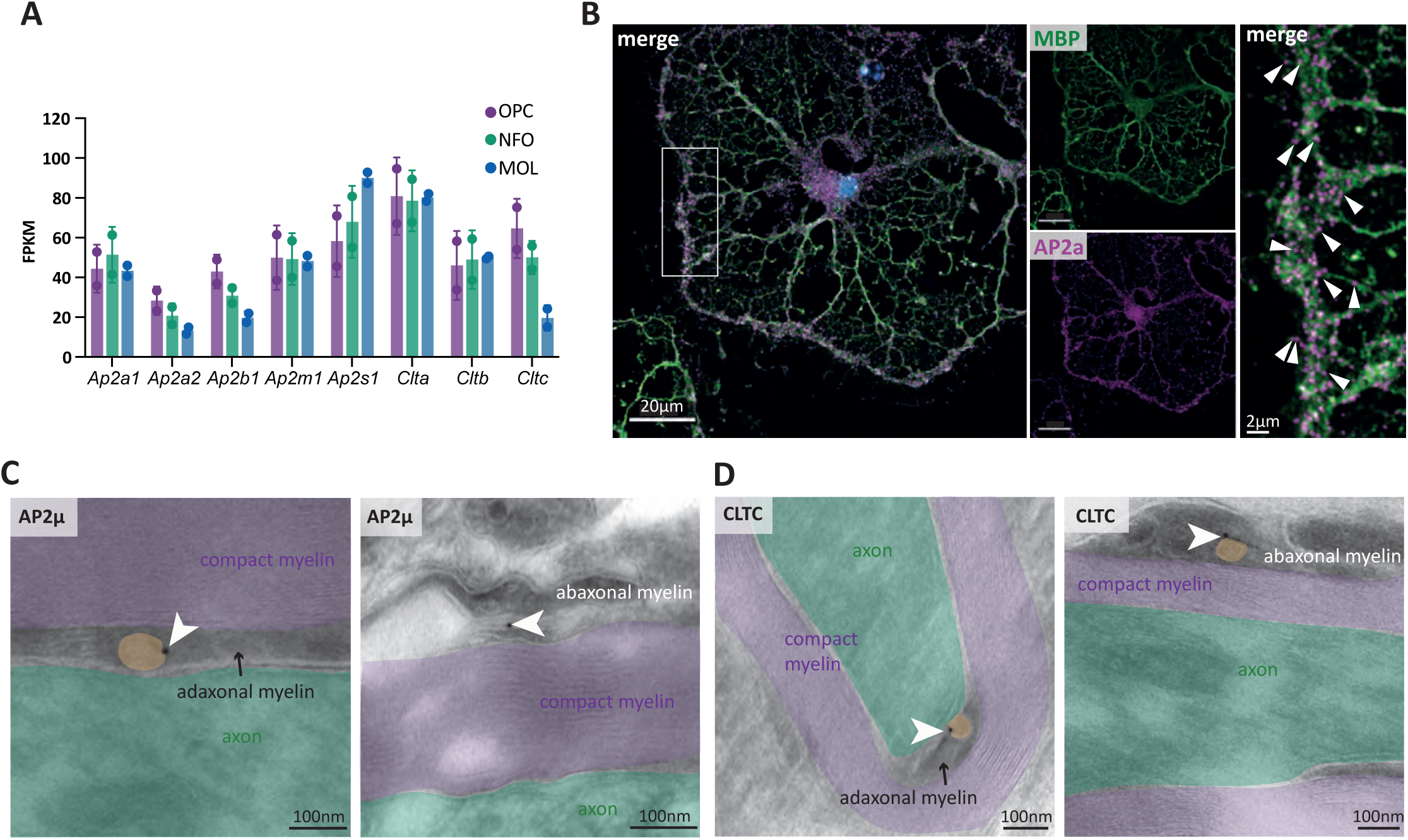
AP2 and clathrin in oligodendrocytes and myelin. **A** Abundance of mRNAs encoding proteins involved in clathrin-mediated endocytosis (CME) replotted from published RNA-Seq data of cells of the oligodendrocyte linage immunopanned from mouse cortex (Zhang et al., 2014). Note that relevant transcripts are expressed in oligodendrocytes. Data shown as mean ±SEM; datapoints indicate n=2 experiments. OPC, oligodendrocyte precursor cells (containing 5% microglial contamination); NFO, newly formed oligodendrocytes; MOL, mature oligodendrocytes; FPKM, fragments per kilobase per million mapped reads. **B** Immunolabelling of primary cultured oligodendrocytes 7 days *in vitro* (7 DIV) with antibodies specific for AP2α (magenta) and MBP (green) indicates localization of AP2α-immunopositive puncta in MBP-immunopositive oligodendrocytes. Nuclei were labelled with DAPI (blue). Image shows one MBP-immunopositive oligodendrocyte representative of 3 cultures. **C, D** Immunodetection of adaptor protein complex 2 subunit µ (AP2µ; 10 nm gold particles; white arrowheads pointing at immunogold in **C**) and clathrin heavy chain (CLTC; **D**) on longitudinal optic nerve cryo-sections of wild type mice at 8 months (mo) of age. Note that immunopositive structures were detected in both the adaxonal and abaxonal myelin layers. Compact myelin is false-colored in purple; axons as identified by cytoskeletal elements are indicated in green; clathrin-coated vesicles are false-colored in orange. Images show n=1 mouse representative of n=3 mice.

We then aimed to visualize AP2 in oligodendrocytes. We used magnetic cell separation (MACS) to isolate OPC, which were cultured and differentiated into mature oligodendrocytes. By immunofluorescence labeling after 7 days *in vitro* (DIV), we identified myelinating oligodendrocytes using antibodies specific for the marker myelin basic protein (MBP), as well as AP2 complexes by detecting the AP2 subunit AP2α (**Fig. 1b**). We found a considerable number of AP2α-immunopositive puncta in MBP-immunopositive cells, implying the presence of AP2 complexes in myelinating oligodendrocytes. We note that AP2α-immunopositive puncta localized to oligodendroglial cell bodies and peripheral processes, the latter probably reflecting an *in vitro*-correlate of myelin membranes.

To determine the localization of AP2 complexes and clathrin in myelin *in vivo*, we used cryo-immuno electron microscopy of optic nerves dissected from wild type mice. Using antibodies specific for the AP2 subunit AP2µ **(Fig. 1c**) or clathrin heavy chain (CLTC; **Fig. 1d**) and protein-A coupled to 10 nm gold particles (visible as black puncta in **Fig. 1c, 1d**), both AP2µ and CLTC labeling was readily detected in association with membranes in both the abaxonal and adaxonal non-compact layers of CNS myelin, mainly vesicular structures.

Together these data indicate that the components of the AP2 complexes and clathrin coats required for AP2-dependent endocytosis are expressed in oligodendrocytes and present in CNS myelin sheaths.

### Hypomyelination and myelin pathology upon *Ap2m* recombination in myelinating oligodendrocytes

To recombine the *Ap2m* gene in oligodendrocytes, we used a previously generated mouse line with an allele in which exons 2 and 3 are flanked by loxP sites (floxed), i.e. *Ap2m^flox^* mice (Kononenko et al., 2014). The *Ap2m* gene encodes subunit AP2µ of the AP2 complex; *Ap2m^flox^*mice were previously used to impair the AP2 complex in neurons (Kononenko et al., 2014), cochlear hair cells (Jung et al., 2015), and astrocytes (López-Hernández et al., 2020). To recombine the *Ap2m^flox^* allele specifically in myelinating oligodendrocytes (**Fig. 2a**), *Ap2m^flox^* mice were interbred with *Cnp^Cre^*mice (Lappe-Siefke et al., 2003). By appropriate interbreedings, we generated *Ap2m^flox/flox^;Cnp^Cre^* mice (also termed *Ap2m* cKO) and *Ap2m^flox/flox^* littermate control mice (also termed Ctrl). To assess if the *Ap2m^flox^* allele is recombined in the white matter of *Ap2m* cKO mice, we used PCR on genomic DNA extracted from optic nerves. We found that the recombined *Ap2m^flox^* allele was readily detected in *Ap2m* cKO mice, but not in Ctrl mice (**Supplemental Figs. S1a, S1b**). We then used qRT-PCR with primers flanking exons 2 and 3 to amplify *Ap2m* mRNA from cDNA synthesized from a white matter tract (corpora callosa). Indeed, transcripts of the recombined *Ap2m^flox^*gene (lacking exons 2 and 3) were readily detected in *Ap2m* cKO compared to Ctrl mice (**Supplemental Figs. S1d, S1e**). To examine if *Ap2m* cKO mice display a reduced abundance of AP2 subunits in myelin, we performed immunoblot analyses of myelin biochemically purified from brains at postnatal day 24 (P24). The abundance of AP2µ was considerably reduced in *Ap2m* cKO compared to Ctrl myelin (**Fig. 2b**). Interestingly, the abundance of AP2α was also reduced in *Ap2m* cKO myelin (**Fig. 2b**), suggesting that the other subunits of the AP2 complex are degraded when AP2µ subunits are unavailable. The fact that AP2 subunits are reduced in - but not entirely absent from - *Ap2m* cKO myelin implies partial recombination, i.e. one or even both of the *Ap2m^flox^* alleles may escape recombination in a subset of oligodendrocytes. When detecting the markers myelin-oligodendrocyte glycoprotein (MOG) and myelin-associated glycoprotein (MAG), we found that the relative abundance of MOG was similar while that of MAG was reduced in *Ap2m* cKO compared to Ctrl myelin at P24. Together, *Ap2m* cKO mice provide a novel model that allows studying the consequences for myelination *in vivo* when the AP2-complex is impaired in oligodendrocytes.

**Fig. 2.**
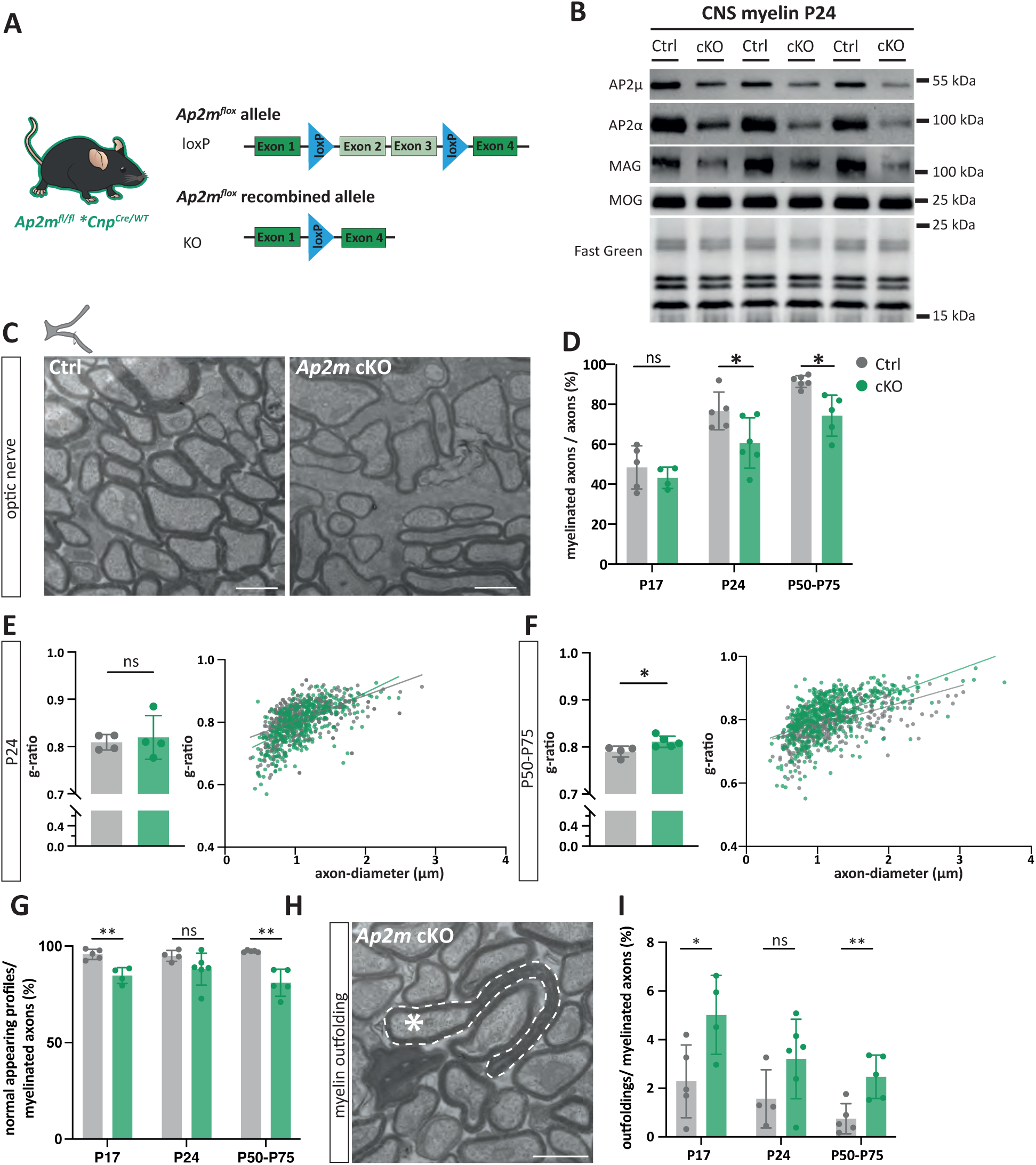
Recombination of *Ap2m* in oligodendrocytes causes hypomyelination and pathological myelin outfoldings. **A** Scheme of the *Ap2m^flox^* allele (Kononenko et al., 2014), in which exons 2 and 3 of the *Ap2m* gene are flanked by loxP-sites to allow *Cre*-mediated recombination. To recombine *Ap2m* in oligodendrocytes, *Ap2m^flox/flox^* (Kononenko et al., 2014) and *Cnp^Cre/WT^* mice (Lappe-Siefke et al., 2003) were interbred. For genomic genotyping PCR and qRT-PCR see **Supplemental data Fig. S1b, S1e**. **B** Immunoblot analysis of myelin purified from brains of *Ap2m^flox/flox^;Cnp^Cre/WT^* (*Ap2m*-cKO) and control (Ctrl) mice at P24 shows a reduced abundance of AP2µ and AP2α in *Ap2m*-cKO myelin. Myelin oligodendrocyte glycoprotein (MOG) was detected as a marker. Fast green protein staining serves as loading and transfer control. Blot shows n=3 biological replicates per genotype. **C** Representative electron micrographs of optic nerve cross sections from Ctrl and *Ap2m*-cKO mice, as exemplified for P50. Scale bar, 1µm. For representative electron micrographs of spinal cord cross sections see **Supplemental data Fig. S2**. **D** Genotype-dependent quantification of electron micrographs as in **C** indicates a reduced percentage of myelinated axons in *Ap2m*-cKO at P24 and P50-P75. Data shown as mean ±SD; datapoints indicate individual mice (n=4-6 per genotype). Two-tailed unpaired t-test with Welch correction, P17 p=0.382, P24 p=0.039, P50-75 p=0.018. **E-F** g-ratio analysis of normal appearing axon-myelin profiles in optic nerves of Ctrl and cKO mice reveals an increased g-ratio indicating thinner myelin sheaths in *Ap2m*-cKO compared to Ctrl mice at P50-P75 (**F**). Bar graphs give mean ±SD; n=4-5 mice per genotype and age; Two-tailed unpaired t-test, P24 p=0.700, P50-70 p=0.033. Scatter plots display g-ratio with respective axon diameters; datapoints show 100-200 axons per mouse from 4-5 mice per genotype and age. **G** Genotype-dependent quantification of electron micrographs as in **C** shows a decreased percentage of myelinated axons that appears non-pathological in *Ap2m*-cKO mice at P17 and P50-P75. Data given as mean ±SD; datapoints indicate individual mice; n=4-6 mice per genotype and age. Two-tailed unpaired t-test with Welch correction, P17 p=0.006, P24 p=0.102, P50-75 p=0.006. **H-I** Electron micrographs of optic nerves showing a representative pathological myelin outfolding in an *Ap2m*-cKO mouse. Stippled white line indicates a myelin outfolding, the asterisk marks the associated axon (**H)**. Scale bar 1µm (**H)**. Genotype-dependent quantification indicates an increased percentage of myelinated axons that display myelin outfoldings in *Ap2m* cKO at P17 and P50-P75 **(I)**. Data shown as mean ±SD; datapoints indicate individual mice; n=4-6 mice per genotype and age. Two-tailed unpaired t-test with Welch correction, P17 p=0.034, P24 p=0.126, P50-75 p=0.008. For assessment of secondary neuropathology see **Supplemental data Figs. S3a-S3d**.

To assess developmental myelination in *Ap2m* cKO mice we used transmission electron microscopy (TEM) of optic nerves **(Fig. 2c)** at three different ages (P17, P24, P50-P75), allowing to identify structural changes of the axon/myelin-unit. By quantitative assessment of electron micrographs, we found a significantly reduced fraction of axons being myelinated at both of the older ages **(Fig. 2d)**, though not at P17. When assessing myelin sheath thickness, we found significantly increased g-ratios reflecting reduced myelin thickness in the optic nerves of *Ap2m* cKO mice at the oldest assessed age **(Fig. 2f)**, though not at P24 **(Fig. 2e)**. Together, these data indicate moderate but significant hypomyelination of axons in *Ap2m* cKO mice, in agreement with the hypothesis that normal developmental CNS myelination by oligodendrocytes involves the AP2-complex.

Surprisingly, when assessing the electron micrographs, we noticed that the fraction of axon/myelin-units that displayed a normal morphology was reduced in *Ap2m* cKO compared to Ctrl optic nerves **(Fig. 2g)**. The most evident pathological feature was a considerable number of axon/myelin-units displaying myelin outfoldings (exemplified in **Fig. 2h**). Myelin outfoldings are large sheets of compacted multimembrane myelin stacks that extend away from the myelinated axon; a frequent type of myelin pathology (Peters, 2002; Patzig et al., 2016; Djannatian et al., 2023; Steyer et al., 2023; Collins et al., 2024) also referred to as focal hypermyelination. Indeed, by quantitative assessment of electron micrographs, the percentage of axon/myelin-units displaying myelin outfoldings was significantly increased in *Ap2m* cKO compared to Ctrl optic nerves at two of the analyzed timepoints **(Fig. 2i)**. At age P24, *Ap2m* cKO mice displayed a trend toward an increased frequency of myelin outfoldings that did not reach significance **(Fig. 2i)**. To assess if myelin outfoldings also emerge in CNS regions other than the optic nerve, we examined the ventral spinal cord at P24 by TEM. Indeed, spinal cords of *Ap2m* cKO mice but not Ctrl mice displayed large myelin outfoldings (see representative images in **Supplemental Fig. S2**).

Together, a combined phenotype of hypomyelination and focal hypermyelination emerges in *Ap2m* cKO mice. It is plausible that the observed hypomyelination reflects diminished developmental biogenesis of myelin when the AP2 complex is impaired in oligodendrocytes. However, *Ap2m* cKO mice, in which the *Ap2m* gene is recombined in myelinating oligodendrocytes already during development, are not suited to assess if the focal hypermyelination is a secondary consequence of impaired developmental myelination or reflects impaired post-developmental maintenance of myelin sheaths. To test these hypotheses, we turned to mice in which the *Ap2m* gene is recombined in oligodendrocytes after myelin biogenesis has been largely completed.

### Recombination of *Ap2m* in oligodendrocytes of adult mice causes progressive myelin pathology

To assess the consequences of impairing the AP2 complex specifically in oligodendrocytes of adult mice, *Ap2m^flox^* mice were interbred with *Plp^CreERT2^* mice (Leone et al. 2003), which express a tamoxifen-inducible variant of Cre recombinase (Feil et al., 1997) in myelinating oligodendrocytes, thus allowing temporal control of *Ap2m^flox^*gene recombination. By appropriate interbreedings, we obtained *Ap2m^flox/flox^; Plp^CreERT2^* mice (also termed *Ap2m* iKO) mice and *Ap2m^flox/flox^* littermate controls (Ctrl). Both *Ap2m* iKO and Ctrl mice were injected with tamoxifen at the age of 8 weeks (**Fig. 3a**). According to genomic PCR, the *Ap2m^flox^* allele was recombined in *Ap2m* iKO but not Ctrl mice (**Supplemental Figs. S1a, S1c**), and by qRT-PCR transcripts of the recombined *Ap2m^flox^*gene (lacking exons 2 and 3) were readily detected in cDNA synthesized from a white matter tract (corpora callosa) in *Ap2m* cKO compared to Ctrl mice (**Supplemental Figs. S1d, S1f**).

**Fig. 3:**
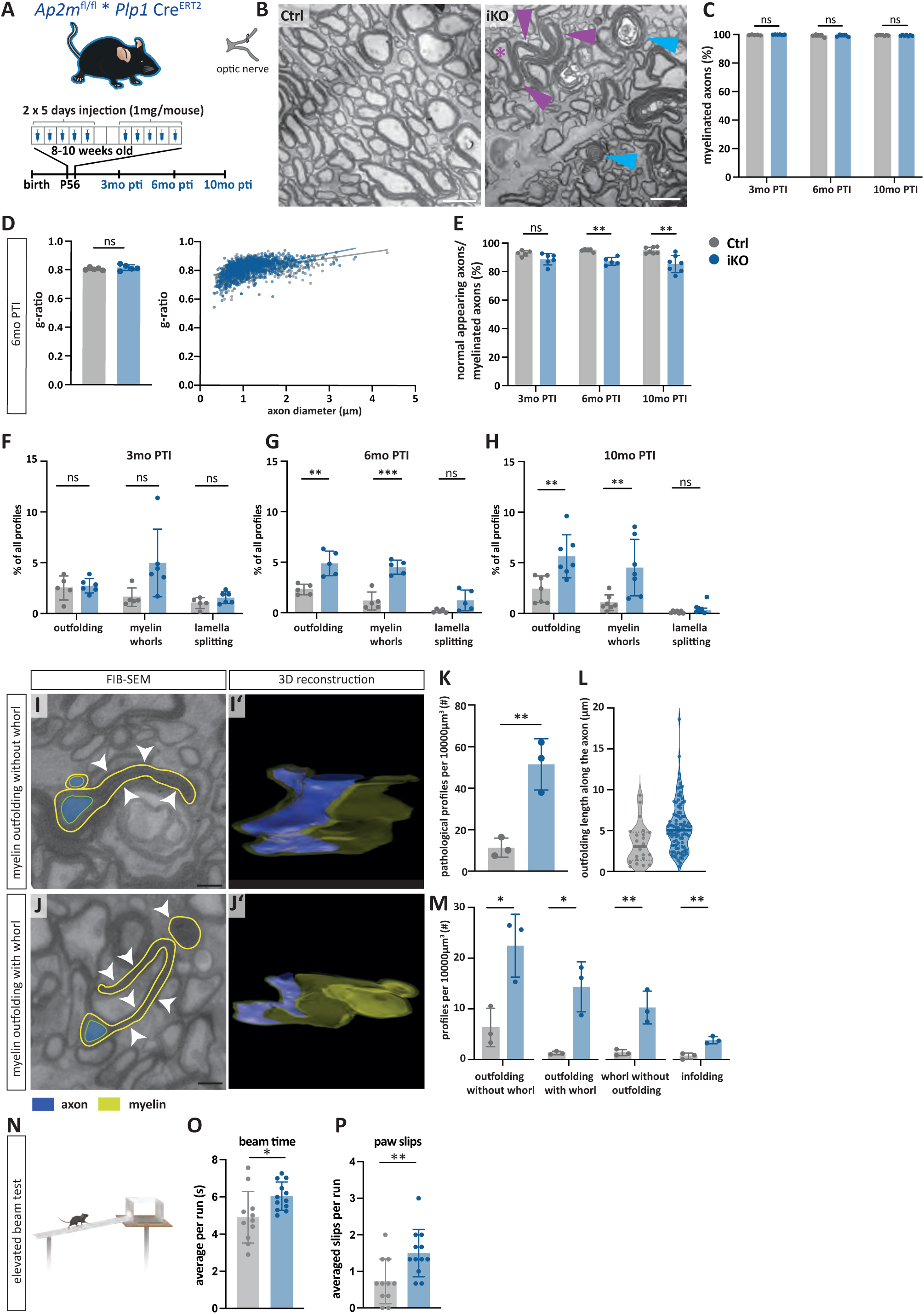
*Ap2m*-recombination in oligodendrocytes of adult mice causes myelin pathology. **A** To recombine *Ap2m* in oligodendrocytes of adult mice, *Ap2m^flox/flox^* (Kononenko et al., 2014) and *Plp-Cre^ERT2^* mice (Leone et al. 2003) were interbred; *Ap2m^flox/flox^;Plp^CreERT2^* (*Ap2m*-iKO) and control (Ctrl) mice were injected with Tamoxifen at 8-10 weeks of age, and analyzed 3, 6 or 10 months (mo) post tamoxifen injection (PTI). For genomic genotyping PCR and qRT-PCR see **Supplemental data Fig. S1c, S1f**. **B** Electron micrographs of optic nerve cross sections from Ctrl and *Ap2m*-iKO mice 10 mo PTI. Magenta arrowheads indicate representative myelin outfoldings with the associated axon marked with an asterisk. Blue arrowheads indicate representative myelin whorls. Scale bar, 1 µm. **C** Genotype-dependent quantification of electron micrographs as in **B** shows similar percentage of myelinated axons in Ctrl and *Ap2m*-iKO mice. Data is given as mean ±SD; datapoints represent individual mice; n=5-6 mice per genotype and age. Two tailed unpaired t-test, 3 mo PTI p=0.156, 6 mo PTI p=0.994, 10 mo PTI p=0.40. **D** g-ratio analysis of normal-appearing axon-myelin profiles in optic nerves of Ctrl and *Ap2m*-iKO mice 6 mo PTI indicates similar myelin thickness. Bar graph gives mean ±SD; datapoints indicate individual mice; n=5 mice per genotype; Unpaired Student’s t-test, p=0.335. Scatter plot displays g-ratios with respective axon diameters. Data shows 100-200 axons per mouse from 5 Ctrl and 5 *Ap2m*-iKO mice. **E** Genotype-dependent quantification of electron micrographs as in **B** reveals a progressive decline in the percentage of normal-appearing axon-myelin profiles in *Ap2m*-iKO mice. Two tailed unpaired t-test with Welch’s correction, 3 mo PTI p=0.0558, 6 mo PTI p=0.0003, 10 mo PTI p=0.0019. **F-H** Genotype-dependent quantification of electron micrographs as in **B** revels the percentage of myelinated axons displaying myelin outfoldings, myelin whorls, or lamella splittings 3 mo PTI (**F**), 6 mo PTI (**G**), and 10 mo PTI (**H**). Note that myelin outfoldings and whorls were the most evident myelin pathology in *Ap2m*-iKO mice 6 mo PTI and 10 mo PTI, while lamella splittings were not a significant feature. Data shown as mean ±SD; two-tailed unpaired t-test; **F** outfoldings p=0.7042, myelin whorls p=0.0574, lamella splittings p=0.1509; **G** outfoldings p=0.0024, myelin whorls p=0.0002, lamella splittings p=0.0523; **H** outfoldings p=0.0047, myelin whorls p=0.0079, lamella splittings p=0.0521. Datapoints indicate individual mice; n=5-6 mice per genotype and age. **I-J** Representative focused ion beam-scanning electron microscopy (FIBSEM) micrographs **(I,J)** and three-dimensional reconstruction **(I‘,J‘)** of myelin pathology in *Ap2m*-iKO optic nerves 6 mo PTI. Myelin is marked in green and the corresponding axon in blue. Scale bar, 1µm. In each tissue block, >800 FIB-SEM sections were scanned, resulting in volume stacks of 40-210 µm with a voxel size of 5 nm x 5 nm x 25 nm. Shown are one representative myelin outfolding without whorl **(I,I‘)** and one myelin outfolding with whorl **(J,J‘)**. Three mice per genotype and one tissue block per mouse were analyzed. For reconstructions of representative myelin outfoldings in one tissue block per genotype, also see **Supplemental Videos 1–2**. **K** Genotype-dependent quantification of all pathological profiles as in **I-J** normalized to a stack volume of 10,000 µm^3^. Data shown as mean ±SD; two-tailed unpaired t-test p=0.006; datapoints represent individual mice; n=3 per genotype. **L** Violin plot displays the longitudinal extent along the axon of all myelin outfoldings in 3D volumes as in **I,J** that were entirely contained in the scanned volume. Median with interquartile ranges; datapoints represent individual myelin outfoldings in one tissue block per mouse and three mice per genotype; Median_Ctrl_=3.41 µm, median*_Ap2m_*_-iKO_=5.45, maximum_Ctrl_=10 µm, maximum*_Ap2m_*_-iKO_=18.6 µm. **M** Genotype-dependent quantification of myelin outfoldings without whorls, outfoldings with whorls, whorls without outfoldings, and myelin infoldings, as in **I-J**, normalized to a stack volume of 10,000 µm^3^. Data shown as mean ±SD; outfolding without whorl p=0.018; outfolding with whorl p=0.010; whorl without outfolding p=0.009; infolding p=0.004; Datapoints represent individual mice; n=3 mice per genotype. For assessment of secondary neuropathology see **Supplemental data Figs. S3e-S3h**. **N-P** Elevated beam test assessing motor capabilities of *Ap2m*-iKO and Ctrl mice 10 mo PTI. Note that *Ap2m*-iKO mice require more time **(O)** and display a higher number of paw slips per run on the beam **(P)**. Data shown as mean ±SD; datapoints represent individual mice; n=11-12 per genotype. Unpaired two-tailed t-test **O** p=0.023, **P** p=0.0077.

When using TEM to assess axon/myelin-units in optic nerves of *Ap2m* iKO and Ctrl mice **(Fig. 3b)**, we found no genotype-dependent differences of the fraction of axons that are myelinated 3 months past tamoxifen injection (3 mo PTI), 6 mo PTI, and 10 mo PTI **(Fig. 3c)**. When assessing myelin thickness 6 mo PTI, *Ap2m* iKO and Ctrl mice displayed similar g-ratios **(Fig 3d)**. Together, these data imply that tamoxifen-induced *Ap2m* iKO mice are not hypomyelinated, at least up to 10 mo PTI. The AP2-complex thus appears dispensable for maintaining normal amounts of myelin after developmental myelination is largely completed.

However, when assessing the electron micrographs, we observed a considerable number of pathological-appearing axon/myelin-profiles in *Ap2m* iKO mice (arrowheads in **Fig. 3b)**. Quantitative evaluation showed a significantly reduced frequency of normal-appearing axon/myelin profiles in *Ap2m* iKO compared to Ctrl optic nerves 6 mo PTI and 10 mo PTI **(Fig. 3e)**. When assessing myelin pathology, the fraction of axon-myelin profiles displaying myelin outfoldings or myelin whorls was significantly increased in *Ap2m* iKO compared to Ctrl mice both 6 mo PTI and 10 PTI **(Fig. 3f, 3g, 3h)**. A trend toward a genotype-dependent difference in the frequency of myelin lamellae splittings 6 mo PTI did not reach significance.

In the electron microscopic view, myelin outfoldings appear as pathological myelin structures, in which compact myelin sheaths extend away from a myelinated axon for a longitudinal extent of several micrometers along the internode (Erwig et al., 2019b; Steyer et al., 2023). The adaxonal myelin membrane remains attached to the axonal surface, the periaxonal space appears unaffected, and entire compact myelin sheaths fold out, in some profiles including some cytoplasm in continuity with that of the adaxonal cytoplasmic myelin layer (**Fig. 2h, 3b**). Myelin whorls are highly disorganized profiles defined by the presence of several layers of myelin membranes but devoid of a recognizable axon. They have thus been suggested to reflect pathological axon/myelin profiles, in which the axon has degenerated while the myelin sheath has remained relatively spared, although disordered (Edgar et al., 2009; Buscham et al., 2022). However, by two-dimensional TEM it is not straightforward to conclude whether myelin whorls are indeed remnants of degenerating axon/myelin profiles or particularly complex myelin outfoldings. For a more detailed morphological characterization of pathological myelin outfoldings and myelin whorls in *Ap2m* iKO mice 6 mo PTI we thus used Focused Ion Beam-Scanning Electron Microscopy (FIB-SEM), which allowed the reconstruction of pathological axon/myelin profiles in three dimensions **(Fig. 3i, 3j; Supplemental Movies 1, 2)**. The frequency of pathological myelin profiles was significantly increased in *Ap2m* iKO compared to Ctrl optic nerves **(Fig. 3k)**, in agreement with the conventional quantitative TEM analysis **(Fig. 3g)**. In addition, both the median and maximum lengths of myelin outfoldings longitudinally along their myelinated axon were considerably increased in *Ap2m* iKO (median=5.45 µm, maximum=18.6 µm) compared to Ctrl optic nerves (median=3.41 µm, maximum=10 µm) (**Fig. 3l**). Interestingly, when assessing myelin outfoldings and myelin whorls after 3-dimensional reconstruction, it was evident that a considerable proportion of pathological features comprises both structures **(Fig. 3m)**, thus indicating highly complex pathology. However, not all myelin outfoldings ran – at a different sectioning level - into a myelin whorl or *vice versa*, indicating that the degree of complexity of myelin pathology in *Ap2m* iKO mice displays considerable variability. Yet, the frequency of occurrence of all three features (outfoldings without whorls, outfoldings with whorls, and whorls without outfoldings) was significantly increased in *Ap2m* iKO compared to Ctrl optic nerves, as was the frequency of myelin infoldings into the myelinated axon **(Fig. 3m)**. FIB-SEM and 3-dimensional reconstruction has thus allowed a more refined assessment of myelin pathology compared to 2D-TEM analysis. Together, our morphological analyses revealed a marked increase of pathological myelin profiles in *Ap2m* iKO mice.

Myelin pathology is frequently associated with secondary neuropathology (Lappe-Siefke et al., 2003; Edgar et al., 2009; Barrette et al., 2013; Lüders et al., 2017). To assess if *Ap2m* cKO and *Ap2m* iKO mice display alterations in oligodendrocyte numbers, astrogliosis, microgliosis, or axonal pathology, we performed immunohistological analyses of a representative white matter tract, the hippocampal fimbria (**Supplemental Fig. S3**). *Ap2m* cKO mice were assessed at P17 and P24; *Ap2m* iKO mice were assessed 3 mo PTI, 6 mo PTI, and 10 mo PTI. When using aspartoacylase (ASPA) as a marker for the cell bodies of myelinating oligodendrocytes, no genotype-dependent differences in oligodendrocyte numbers were observed in *Ap2m* cKO (**Supplemental Fig. S3a**) or *Ap2m* iKO fimbriae (**Supplemental Fig. S3e**). When using GFAP as a marker for astrocytes, the area of immunopositivity was unchanged in *Ap2m* cKO compared to Ctrl fimbriae, as well as in *Ap2m* iKO fimbriae 3 mo PTI and 10 mo PTI (**Supplemental Figs. S3b, S3f**). A moderate increase in the area of GFAP-immunopositivity in *Ap2m* iKO compared to Ctrl fimbriae 6 mo PTI reached significance (**Supplemental Fig. S3f**). When using IBA1 as a marker for microglia, the area of immunopositivity was unchanged in *Ap2m* cKO compared to Ctrl fimbriae. However, the area of IBA1-immunopositivity was significantly increased in *Ap2m* iKO compared to Ctrl fimbriae at all three timepoints of analysis (**Supplemental Figs. S3c, S3g**). When immunolabeling amyloid precursor protein (APP) as a marker for pathological axonal swellings, we did not detect genotype-dependent differences in *Ap2m* cKO or *Ap2m* iKO compared to respective Ctrl fimbriae (**Supplemental Fig. S3d**). Together, neuropathological assessment did not reveal signs of secondary neuropathology in *Ap2m* cKO mice; however, signs of a mild microgliosis emerged upon recombination of *Ap2m* in oligodendrocytes of adult mice.

To explore the possibility that motor capabilities are impaired in *Ap2m* iKO mice, we assessed their performance 10 mo PTI using the elevated beam test (Luong et al., 2011). In this experiment, a mouse is placed on an elevated beam **(Fig. 3n)**, and the time and the number of paw slips are measured until the mouse reaches a shelter on a platform at the end of the beam. The task requires precise paw placement and body balance. Compared to Ctrl mice, *Ap2m* iKO mice needed significantly more time to reach the platform and displayed a significantly higher number of paw slips until reaching the platform **(Fig. 3o, 3p)**. These results indicate that motor capabilities are moderately impaired when *Ap2m* is recombined in cells expressing Cre^ERT2^ under control of the *Plp*-promoter in adult mice.

### Lysosomal membrane proteins accumulate in the abaxonal myelin compartment when *Ap2m* is recombined in oligodendrocytes

Next we tested if recombination of *Ap2m* in oligodendrocytes affects the protein composition of myelin. By label-free mass spectrometry, the relative abundance of most of the 686 proteins quantified in myelin biochemically purified from the brains of *Ap2m* cKO mice at P24 was unaltered compared to Ctrl **(Fig. 4a; Supplemental Fig. S4; Supplemental data Table S1)**, including that of selected myelin markers (**Supplemental Fig. S4a**), proteins at the myelin/axon-interface (**Supplemental Fig. S4d**), and other proteins mass spectrometrically identified in myelin such as receptors and receptor-associated proteins the internalization of which could potentially affect myelination (**Supplemental Fig. S4e**). Not surprisingly, the abundance of the subunits of the AP2 complex was reduced in *Ap2m* cKO myelin **(Fig. 4a; Supplemental Fig. S4b)**, in agreement with immunoblot data for AP2µ and AP2α **(Fig. 2b)**. The relative abundance of MAG was moderately but significantly reduced in *Ap2m* cKO compared to Ctrl myelin **(Fig. 4a)**, consistent with immunoblot data **(Fig. 2b)**. Most strikingly, however, the abundance of both lysosome-associated membrane glycoproteins 1 and 2 (LAMP1, LAMP2) was markedly increased in *Ap2m* cKO compared to Ctrl myelin **(Fig. 4a; Supplemental Fig. S4c; Supplemental data Table S1)**, a finding that we were able to validate by immunoblotting **(Fig. 4b)**. Likewise, we used immunoblotting to test if LAMP1 and LAMP2 are also enriched in *Ap2m* iKO compared to Ctrl myelin, which was indeed the case **(Fig. 4c)**. For comparison, the abundance of lysosomal lumen/matrix protein cathepsin D (CTSD) was not increased in *Ap2m* cKO myelin according to immunoblotting **(Fig. 4b)** and proteome analysis **(Supplemental Fig. S4c; Supplemental data Table S1)**, or in *Ap2m* iKO myelin according to immunoblotting **(Fig. 4c)**, compared to respective controls. The relative abundance of MOG and MAG did not differ between *Ap2m* iKO and Ctrl myelin **(Fig. 4c)**. Together, the abundance of LAMP1 and LAMP2 in myelin is highly increased in oligodendroglial *Ap2m* mutants.

**Fig. 4.**
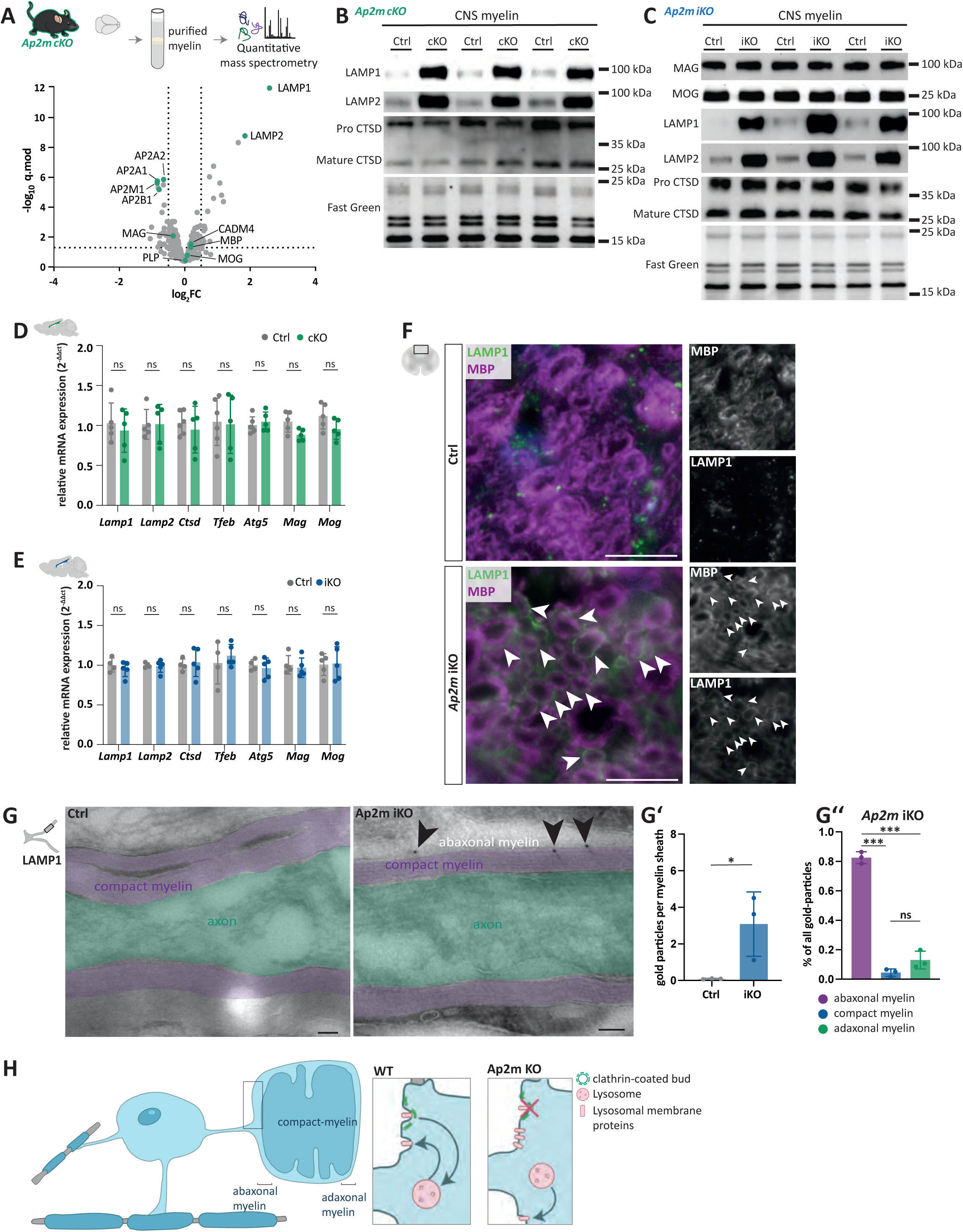
Lysosomal membrane proteins accumulate in abaxonal myelin upon *Ap2m* recombination in oligodendrocytes. **A** Differential quantitative proteome analysis of myelin purified from brains of *Ap2m*-cKO mice. Volcano plot comparing the abundance of proteins in *Ap2m*-cKO and Ctrl myelin at age P24, plotted as the log_2_-transformed fold-change (FC) on the x-axis against the -log_10_-transformed q-value on the y-axis. Stippled lines respectively indicate a log_2_FC of 0.5 and -0.5 and a -log_10_-transformed q-value of 1.301 as significance threshold. Mass spectrometric quantification was performed using 3 biological replicates per genotype with two technical replicates each. Note the strong increase in the abundance of lysosomal-associated membrane proteins (LAMP1, LAMP2) in *Ap2m*-cKO myelin. For heatmap visualization of selected proteins see Supplemental **Figure S4**; for entire dataset see **Supplementary Table 1. B,C** Immunoblot-analysis validates the strong increase in the abundance of LAMP1 and LAMP2 in myelin purified from the brains of *Ap2m*-cKO mice at P24 (**B**), and shows a similar increase in myelin purified from the brains of *Ap2m*-iKO mice compared to respective controls 6 mo PTI (**C**). Cathepsin D (CTSD) was tested as a marker for the lysosomal lumen. Fast green protein staining serves as loading and transfer control. Blots show n=3 per genotype. **D,E** qRT-PCR to determine the abundance of the indicated mRNAs encoding lysosomal and autophagosome proteins in the white matter (corpus callosum) of *Ap2m*-cKO and Ctrl mice at P17 **(D)** and of *Ap2m*-iKO and Ctrl mice 3 mo PTI **(E)**. mRNA abundance was normalized to *Rps13* and *Rplp0* as housekeepers. Note that the abundance of all tested mRNAs was unaltered in *Ap2m*-mutant mice. Data are shown as mean ±SD; datapoints represent individual mice; n=4-6; not significant according to two-tailed unpaired t-test **D** *Lamp1* p=0.887, *Lamp2* p=0.666, *Ctsd* p=0.543, *Tfeb* p=0.977, *Mag* p=0.105, *Mog* p=0.536; **E** *Lamp1* p= 0.436, *Lamp2* p= 0.775, *Ctsd* p=0.734, *Tfeb* p=0.5278, *Atg5* p= 0.608, *Mag* p=0.639, *Mog* p=0.907. For assessment of autophagy markers by immunoblot and immunohistochemistry see **Supplemental data Fig. S4**. **F** Immunohistochemical analysis of LAMP1 (green) and the myelin marker MBP (magenta) in spinal cord cross sections from Ctrl (top) and *Ap2m*-iKO (bottom) mice. Note that LAMP1 was readily detected in myelin in the white matter of *Ap2m*-iKO mice. Scale bar, 10µm. **G** Immunodetection of LAMP1 (10 nm gold particles; black arrowheads pointing at black immunogold) on longitudinal optic nerve cryo-sections of *Ap2m*-iKO and Ctrl mice 6 mo PTI. Compact myelin is false-colored in purple and the axon is marked in green. Note that LAMP1-immunogold was preferentially detected in the abaxonal myelin layer of *Ap2m*-iKO mice. Images show n=1 mouse representative of n=3 mice per genotype. Scale bar, 100 nm. **G‘-G‘‘** Genotype-dependent quantification on images as in **G** of LAMP1 immunogold particles in myelin profiles (**G‘**), and subcompartment-dependent quantification of LAMP1 immunogold in abaxonal, compact, and adaxonal myelin of *Ap2m*-iKO mice (**G‘‘**). Data given as mean ±SD; datapoints represent individual mice; n=3. **G‘** unpaired Student’s t-test p=0.026; **G‘‘** 2-way-ANOVA abaxonal vs. compact myelin p<0.001, abaxonal myelin vs. adaxonal myelin p<0.001, compact myelin vs. adaxonal myelin p=0.247. Note that LAMP1 immunogold was virtually undetectable in myelin of Ctrl mice and strongly enriched in the abaxonal myelin layer of *Ap2m*-iKO mice. **H** Hypothetical model of the endocytotic retrieval of lysosomal membrane proteins from abaxonal myelin membranes in wild-type (Ctrl) oligodendrocytes, and impaired retrieval in *Ap2m*-mutant (*Ap2m* KO) oligodendrocytes.

When *Ap2m* is recombined in astrocytes (López-Hernández et al., 2020), the emerging increased abundance of lysosomal proteins coincided with transcriptional activation of lysosome-related genes. We thus tested if the relative abundance of *Lamp1* and *Lamp2* transcripts is also increased upon oligodendroglial recombination of *Ap2m.* However, qRT-PCR analysis of cDNA generated from a model white matter tract (the corpus callosum) dissected from *Ap2m* cKO or iKO and respective Ctrl mice, did not detect genotype-dependent differences **(Fig. 4d, 4e)**. Likewise, the relative abundance of transcripts encoding the lysosomal marker CTSD, autophagy-related transcription factor EB (TFEB), autophagy-related protein 5 (ATG5), MAG or MOG also did not differ between the genotypes **(Fig. 4d, 4e)**. Together, this implies that the increased abundance of LAMP1 and LAMP2 in *Ap2m* cKO and iKO myelin is due to a post-transcriptional effect, in difference to the transcriptional changes observed upon recombination of *Ap2m* in astrocytes (López-Hernández et al., 2020).

Given that our data imply that autophagy may not be activated when *Ap2m* is recombined in oligodendrocytes, we tested for additional autophagosomal markers (Klionsky et al., 2021). By immunoblotting, we found no genotype-dependent difference in the relative abundance of sequestosome 1 (P62) or microtubule-associated proteins 1A/1B light chain 3A (LC3-I, LC3-II) in brain lysates or myelin purified from *Ap2m* cKO or *Ap2m* iKO compared to respective control mice (**Supplemental Fig. S5a-S5d**). By immunofluorescence labeling, we found no indication of an altered abundance or localization of ATG5 or TFEB in spinal cord cross sections of *Ap2m* iKO compared to Ctrl mice (**Supplemental Fig. S5e, S5f**), in agreement with the qRT-PCR data **(Fig. 4d, 4e)**. Together, these experiments show no evidence of activation of autophagy when *Ap2m* is recombined in oligodendrocytes.

To assess the localization of LAMP1 in myelin upon oligodendroglial recombination of *Ap2m*, we first performed immunofluorescence labeling experiments using cross sections of spinal cords dissected from *Ap2m* iKO and Ctrl mice 6 mo PTI. The immunofluorescence signal of LAMP1 (green in **Fig. 4f**) was strongly increased in the white matter of *Ap2m* iKO spinal cords, in proximity to but not overlapping with MBP-immunolabelling (magenta in **Fig. 4f**) as a marker for compact myelin. These data indicate that LAMP1-immunolabeling in *Ap2m* iKO myelin is most prominent in the non-compacted abaxonal myelin layer. To further specify the precise localization of LAMP1, we used cryo-immuno electron microscopy to detect LAMP1 on longitudinal cryosections of optic nerves dissected from *Ap2m* iKO and Ctrl mice **(Fig. 4g)**. By quantitative assessment, LAMP1 immunogold labelling was virtually undetectable in myelin of Ctrl mice **(Fig. 4g‘).** Conversely LAMP1 immunogold labelling was readily detected in myelin profiles of *Ap2m* iKO mice **(Fig. 4g‘).** Here, LAMP1 immunogold labelling was strongly enriched in the abaxonal myelin layer compared to compact myelin or the adaxonal layer **(Fig. 4g’’)**. In this experiment, we did not observe LAMP1 immunogold labelling associated with vesicular structures in the myelin sheath. Together, and the fact notwithstanding that autophagy is not induced, these data show that lysosomal membrane proteins accumulate in the abaxonal myelin layer when *Ap2m* is recombined in oligodendrocytes **(Fig. 4h)**.

## Discussion

Here we used mouse genetics to test if developmental myelin biogenesis *in vivo* involves the AP2 complex that mediates AP2-dependent endocytosis (Mettlen et al., 2018; Azarnia Tehran et al., 2019; Chen and Schmid, 2020; López-Hernández et al., 2020; Smith and Smith, 2022). This hypothesis was supported by the finding that a reduced percentage of axons is myelinated and myelin thickness is reduced upon recombination of the *Ap2m*-gene in *Cnp*-expressing myelinating oligodendrocytes. Unexpectedly, pathological focal hypermyelination was also a feature in *Ap2m* cKO mice, which we hypothesized to reflect impaired myelin maintenance rather than biogenesis. To test this notion, we recombined the *Ap2m*-gene in *Plp*-expressing myelinating oligodendrocytes specifically in adult mice after developmental myelination is largely completed. *Ap2m* iKO mice displayed normal developmental myelination but late-onset progressive formation of pathological myelin outfoldings and whorls. Our data imply that oligodendrocytes cell-autonomously contribute to the long-term maintenance of their myelin sheaths involving AP2-dependent endocytosis.

### The AP2 complex facilitates myelin biogenesis

According to theoretical calculations (Pfeiffer et al., 1993), each oligodendrocyte synthesizes up to 5000 µm^2^ of myelin membrane per day during active myelination. At the same time, the intracellular transport and sorting of myelin constituents must be spatio-temporally coordinated to facilitate the specialized morphology and molecular composition of healthy myelin sheaths (Hildebrand et al., 1993; Pfeiffer et al., 1993). For example, the glycolipid galactosylceramide (GalC) is transferred into myelin membranes via contact sites between endoplasmic reticulum and plasma membrane (Wu et al., 2024), and some myelin proteins are locally translated from transcripts at free ribosomes localized to myelin (Thakurela et al., 2016; Yergert et al., 2021), including MBP (Ainger et al., 1993; Müller et al., 2013). However, most myelin components are considered to be trafficked via vesicles before their integration into the growing sheath via exocytosis (Trajkovic et al., 2006; Winterstein et al., 2008; Feldmann et al., 2009, 2011). When vesicular SNARE proteins (VAMP2, VAMP3) are cleaved by Botulinum neurotoxin B and thus inactivated in oligodendrocytes of *iBot;Cnp^Cre^*mice, the fusion of vesicles with the oligodendroglial plasma membrane is blocked (Lam et al., 2022). *iBot;Cnp^Cre^* mice display pronounced developmental hypomyelination, confirming that myelin membrane expansion involves exocytic fusion of vesicles comprising myelin constituents (Lam et al., 2022).

In multiple cell types, including neurons, epithelial cells, and endothelial cells, the delivery of membrane to the destined cell surface domain involves transcytosis as a major transport route (Mostov and Cardone, 1995; Hémar et al., 1997; Middleton et al., 1997; Ribeiro et al., 2019). According to this concept, newly synthesized membrane components fuse with the plasma membrane via exocytosis of Golgi-derived vesicles, then undergo endocytic uptake from the plasma membrane and are trafficked into the endosomal/lysosomal system. From there, they are subsequently recycled, i.e. integrated into the plasma membrane, thereby facilitating the molecular specialization of cell surface domains. In analogy, we followed the hypothesis that myelin constituents may undergo endocytosis before their eventual integration into the growing sheath *in vivo*. This hypothesis had been supported by prior experiments using cultured oligodendrocytes, which indicated that the most abundant transmembrane myelin proteins undergo endocytosis at least *in vitro*, i.e., proteolipid protein (PLP), myelin-associated glycoprotein (MAG), and myelin-oligodendrocyte glycoprotein (MOG). Interestingly, endocytosis of PLP and its subsequent trafficking into myelin-like membranes is independent of clathrin (Trajkovic et al., 2006; Winterstein et al., 2008; Baron et al., 2015), while MAG and MOG are endocytosed via clathrin-dependent mechanisms (Winterstein et al., 2008). Together, these *in vitro* data are consistent with the hypothesis that myelin biogenesis involves endocytic recycling (Winterstein et al., 2008). It has remained unknown, however, if endocytic mechanisms are indeed required for normal myelination *in vivo*, and, if so, for bulk trafficking of myelin membranes or for particular myelin proteins.

We thus hypothesized that endocytosis is required for normal myelin formation *in vivo*. For gene targeting, we selected the *Ap2m* gene encoding the µ-subunit of the AP2 complex because it is essential for formation and function of the AP2 complex, which mediates assembly of clathrin-coated vesicles at the plasma membrane (Mettlen et al., 2018; Azarnia Tehran et al., 2019; Chen and Schmid, 2020; López-Hernández et al., 2020; Smith and Smith, 2022). Our data show that recombination of the *Ap2m* gene in myelinating oligodendrocytes expressing Cre recombinase under control of the *Cnp*-promoter (Lappe-Siefke et al., 2003) display developmental hypomyelination, as indicated by a reduced percentage of axons being myelinated, and thinner myelin sheaths. Thus, the AP2 complex facilitates normal myelin formation *in vivo*. We note, however, that the formation of myelin sheaths is not entirely abolished in *Ap2m* cKO mice, suggesting that non-AP2-dependent trafficking routes also allow bulk trafficking of future myelin membrane into the growing sheath, though less efficiently.

Our mass spectrometric analysis of biochemically purified myelin indicates that *Ap2m* recombination in oligodendrocytes leads to distinct changes in the molecular composition of myelin. Among the canonical myelin proteins, this solely affects MAG. The abundance of MAG in *Ap2m* cKO myelin is moderately but significantly reduced, probably reflecting the fact that its trafficking in oligodendrocytes during myelin biogenesis involves AP2-dependent endocytosis *in vivo*, as suggested by previous *in vitro* analyses (Winterstein et al., 2008). Indeed, the cytoplasmic domain of the large splice isoform of MAG (L-MAG) comprises amino acid motifs that serve as endocytosis signals (Bö et al., 1995). We note that the abundance of MAG in myelin is unchanged when *Ap2m* is recombined in oligodendrocytes of adult mice (*Ap2m* iKO). This probably reflects that their myelin sheaths are initially formed normally, including a normal abundance of MAG, and that subsequent myelin turnover in adult mice is very slow as it measures in months (Fischer and Morell, 1974; Fornasiero et al., 2018; Lüders et al., 2019), allowing for the possibility that AP2-independent routes facilitate trafficking of a number of MAG molecules into myelin sufficient to replace degraded MAG, leading to normal-appearing abundance of MAG in myelin of *Ap2m* iKO mice. Given that *Mag^null/null^* mice display a reduced percentage of myelinated axons (Bartsch et al., 1997; Li et al., 1998; Biffiger et al., 2000; Pernet et al., 2008; Patzig et al., 2016; Steyer et al., 2023), it is tempting to speculate that the reduced abundance of MAG in *Ap2m* cKO myelin may also contribute to the hypomyelinating phenotype of *Ap2m* cKO mice. However, a direct comparison is not trivial because the relative abundance of MAG in *Ap2m* mutant myelin is only moderately reduced, while *Mag^null/null^* mice entirely lack MAG expression.

On the other hand, except for MAG, the abundance of all other canonical myelin proteins is not changed in *Ap2m* cKO mice, including PLP, MBP, and MOG. This finding was expected for PLP, the endocytosis of which is independent of clathrin in cultured oligodendrocytes (Trajkovic et al., 2006; Winterstein et al., 2008; Baron et al., 2015), and for MBP, which is not a transmembrane protein and incorporated into myelin independent of vesicular transport (Ainger et al., 1993; Müller et al., 2013). Yet, the relative abundance of MOG in myelin of *Ap2m* cKO mice was unchanged although MOG is endocytosed via a clathrin-dependent mechanism in cultured oligodendrocytes (Winterstein et al., 2008). Notably, according to cryo immuno-electron microscopy, MOG localizes to the abaxonal myelin surface (Brunner et al., 1989) while MAG is enriched in the adaxonal layer (Trapp et al., 1989). It is possible that intracellular transport of MAG towards the growing myelin sheath during development involves AP2-dependent endocytosis, while trafficking of MOG to its normal localization in abaxonal myelin is independent of AP2-dependent endocytosis *in vivo*.

### Abaxonal myelin as a site for retrieving myelin constituents

How can proteins or membrane patches be retrieved from the compactly layered myelin sheath? It is difficult to conceive how vesicles could bud off from membranes within densely compacted myelin layers. In the noncompacted myelin compartments, however, intracellular surfaces of myelin membranes are flanked by cytoplasm, including in the adaxonal and abaxonal myelin layers. These cytosolic channels provide cytoplasmic continuity with the oligodendroglial cell body and comprise cellular organelles and vesicles (Snaidero and Simons, 2017; Frühbeis et al., 2020; Trevisiol et al., 2020; Chapple et al., 2024). Indeed, adaxonal and abaxonal myelin exhibit AP2- and clathrin-immunopositive membrane buds and vesicles, indicating that non-compact myelin contains the molecular machinery to mediate AP2-dependent endocytosis, including that of LAMP1/2.

The most striking change of the molecular composition of myelin in both *Ap2m* cKO and iKO mice is the marked enrichment of the lysosomal membrane proteins LAMP1/2 in abaxonal myelin. This implies that, under normal conditions, lysosomal membrane proteins are integrated into but do not accumulate in the abaxonal myelin membrane. Normal intracellular trafficking of LAMP1/2 in oligodendrocytes is thus likely to involve AP2-dependent endocytosis at this membrane. Indeed, both LAMP1 and LAMP2 contain tyrosine-based sequence motifs for endocytosis via clathrin-coated vesicles (Bonifacino and Traub, 2003). The occurrence of LAMP1/2 in abaxonal myelin may simply reflect that after biosynthesis they are trafficked to lysosomes via the abaxonal myelin membrane (Janvier and Bonifacino, 2005; López-Hernández et al., 2020). Yet, given that myelin biogenesis comprises a step of LAMP1/2-containing endosomes/lysosomes undergoing exocytosis to deliver myelin constituents (Trajkovic et al., 2006; Feldmann et al., 2011; Shen et al., 2016), it is plausible that LAMP1/2 are incorporated into myelin as a side-product of normal lysosomal/endosomal exocytosis, but accumulate when their AP2-dependent retrieval is impaired.

We note that, in neurons and cochlear hair cells, the AP2 complex has also been implicated in the recycling of synaptic vesicles for their ultrafast exocytosis (Kononenko et al., 2014; Jung et al., 2015). Notably, synaptic exocytosis requires prompt replenishment of vesicles whereas myelin turnover measures in months (Fischer and Morell, 1974; Fornasiero et al., 2018; Lüders et al., 2019). Yet, we can not formally exclude that exocytosis of endosomes/lysosomes is also impaired in oligodendroglial *Ap2m* mutants. However, it is plausible that endocytic membrane retrieval from abaxonal myelin is predominantly affected when considering that the abundance of LAMP1/LAMP2 but not of a marker for the lysosomal lumen (CTSD) or other endo/lysosomal markers is increased, and that LAMP1 accumulates in the abaxonal membrane but not in vesicular or endo/lysosomal structures, in myelin of oligodendroglial *Ap2m* mutants.

While adaxonal myelin is considered the preferential site for the integration of newly synthesized myelin constituents at least during active myelination (Snaidero et al., 2014; Snaidero and Simons, 2017; Meschkat et al., 2022), the abaxonal layer thus emerges as an active site for retrieving membrane constituents from myelin, thereby providing spatial segregation of these functions. While it is probable that the aberrant enrichment of LAMP1/2 in the abaxonal myelin layer reflects impaired retrieval from this membrane, we cannot formally exclude that it secondarily contributes to the observed myelin pathology.

Together, our findings indicate that the AP2 complex is involved in the biogenesis and maintenance of healthy myelin sheaths. On the other hand, the observation that *Ap2m* cKO mice are hypomyelinated but not entirely dysmyelinated implies that the AP2 complex does facilitate trafficking of myelin membranes to some extent, but that AP2-independent trafficking routes also contribute, allowing partial myelination when *Ap2m* is recombined in oligodendrocytes.

### Maintenance of myelin structure

Once myelin sheaths have developmentally formed, the continued biosynthesis of myelin membranes throughout adulthood (Yeung et al., 2014; Baraban et al., 2016; Lüders et al., 2017; Meschkat et al., 2022) requires the removal of aged myelin. Patches of myelin sheaths have been shown to be removed via extrinsic degradation of myelin membranes by phagocytic cells, requiring fragmentation of myelin sheaths or their shedding from the axon. Indeed, myelin can be phagocytosed by microglia or astrocytes during normal development and in pathological conditions (Neumann et al., 2009; Lampron et al., 2015; Safaiyan et al., 2016; Ponath et al., 2017; Hughes and Appel, 2020; Sen et al., 2022; Wan et al., 2022b; Djannatian et al., 2023). However, recent analyses of *Atg5*-cKO and *Atg7*-cKO mice, in which macroautophagy in oligodendrocytes is affected, led to the notion that oligodendrocytes may cell-autonomously contribute to degrading and remodeling myelin membranes (Aber et al., 2022; Zhang et al., 2023; Chen et al., 2024; Ktena et al.). In agreement with this notion, genetic impairment of autophagy in oligodendrocytes results in a late-onset, progressive increase in both myelin thickness and pathological myelin structure, including myelin outfoldings (Aber et al., 2022; Zhang et al., 2023). The emergence of myelin outfoldings and whorls 6 months after tamoxifen-induced recombination of *Ap2m* in oligodendrocytes supports the view that constant removal of aged myelin components is crucial for the long-term integrity of healthy axon/myelin profiles. It is thus plausible that AP2-dependent mechanisms facilitate removal of pathological myelin in an oligodendrocyte-autonomous manner, in addition to microglial and astrocytic phagocytosis of myelin. Yet, considering that *Ap2m* mutants neither displayed increased myelin thickness nor showed evidence of induction of autophagy, AP2-dependent mechanisms of maintaining myelin structure are likely to differ from those reported for *Atg5*-cKO and *Atg7*-cKO mice.

Myelin outfoldings observed during development have been interpreted as a reservoir that allows extending the myelin sheath during longitudinal growth of the myelinated axon (Cullen and Def. Webster, 1979; Snaidero et al., 2014; Djannatian et al., 2023). Several studies have related the emergence of pathological myelin outfoldings to a dysregulation of the actin cytoskeleton causing uncontrolled growth of myelin sheaths after longitudinal axon growth has terminated in the adult CNS (Katanov et al., 2020; Iyer et al., 2024). On the other hand, myelin outfoldings are a major pathology that emerges when myelin stability is impaired in mice lacking the septin filaments that scaffold mature CNS myelin (Patzig et al., 2016; Erwig et al., 2019b) or the myelin adhesion proteins CADM4 (Elazar et al., 2019) or MAG (Li et al., 1994, 1998; Biffiger et al., 2000; Steyer et al., 2023). Considering that myelin proteome analyses revealed a moderately but significantly reduced abundance of MAG in *Ap2m* cKO myelin - but no changes affecting the abundance of actin-related proteins, septins, or CADM4 - the reduced abundance of MAG may contribute to the emergence of myelin outfoldings of *Ap2m* cKO mice. Accordingly, *Mag^null/null^* mice display myelin outfoldings (Li et al., 1994, 1998; Biffiger et al., 2000; Patzig et al., 2016; Steyer et al., 2023). However, *Mag^null/null^* mice entirely lack MAG, while its abundance in *Ap2m* cKO myelin is just moderately reduced, making direct comparisons difficult. Moreover, the abundance of MAG is not reduced in myelin of *Ap2m* iKO mice, which also display myelin outfoldings. In summary, we can not formally exclude that impaired myelin stability may contribute to the emergence of myelin outfoldings in *Ap2m* mutants; however it is more plausible that impaired retrieval of myelin constituents causes disproportionate growth of sheaths.

### Limitations of the study

While it is plausible that the accumulation of lysosomal membrane proteins in abaxonal myelin and the emergence of myelin outfoldings are independent consequences of recombining *Ap2m* in oligodendrocytes, we cannot formally exclude that accumulation of LAMP1/2 in abaxonal myelin secondarily induces the emergence of myelin outfoldings. It will thus be an interesting task for future experiments to generate transgenic mice suited to test if forced overexpression of LAMP1/2 specifically in oligodendrocytes leads to their accumulation in abaxonal myelin, and if so, whether this causes the emergence of myelin outfoldings.

Our analysis focuses on the cell-autonomous relevance of the AP2 complex in myelinating oligodendrocytes for the molecular composition and structure of CNS myelin. It will be a relevant future task to explore the possibility of an interplay between oligodendrocyte-autonomous mechanisms and pruning by astrocytes or microglia in maintaining myelin sheaths. For example, intercellular signalling may link oligodendroglial membrane retrieval to phagocytosis by microglia or astrocytes via cell surface or secreted ligands and surface receptors on the respective other cell type, thereby regulating myelin homeostasis in the mature, ageing, or pathological nervous system. Accumulation and exposure of LAMP1/2 on the abaxonal myelin surface may be a potential candidate process for myelin-to-astrocyte/microglia signalling. However, its pathobiological relevance remains to be tested, as well as in which myelin-related disorders this phenomenon occurs beyond *Ap2m* mutant mice.

## Conclusion

This work shows that the AP2 complex, which facilitates AP2-dependent endocytosis, in oligodendrocytes is required for normal biogenesis, molecular composition, and structure of myelin, including the prevention of the emergence of focal hypermyelination. The abaxonal myelin layer is revealed as an active site for AP2-dependent retrieval of membrane constituents from the myelin sheath. Importantly, our data support the emerging concept that, in addition to astrocytes and microglia, oligodendrocytes cell-autonomously contribute to the maintenance of mature sheaths, via an AP2-dependent retrieval mechanism.

## Methods

### Mouse lines

Mice harboring an allele of the *Ap2m* gene in which exons 2 and 3 are flanked by loxP sites (*Ap2m^flox^*allele) (Kononenko et al., 2014) were imported from the Leibniz Forschungsinstitut für Molekulare Pharmakologie (FMP, Berlin, Germany). To recombine *Ap2m* in myelinating oligodendrocytes, *Ap2m^flox^* mice were interbred with mice expressing *Cre* recombinase under control of the *Cnp* promoter (*Cnp^Cre^*) (Lappe-Siefke et al., 2003), yielding *Ap2m^flox/flox^;Cnp^Cre^*mice (also termed *Ap2m* cKO) and *Ap2m^flox/flox^* littermate control mice. To recombine *Ap2m* in myelinating oligodendrocytes of adult mice, *Ap2m^flox^* mice were interbred with mice expressing tamoxifen-inducible Cre recombinase (Feil et al., 1997) under control of the *Plp* promotor (*Plp^CreERT2^* mice) (Leone et al., 2003), yielding *Ap2m^flox/flox^; Plp^CreERT2^* (also termed *Ap2m* iKO) and *Ap2m^flox/flox^* littermate control (Ctrl) mice. At the age of 8 weeks, male *Ap2m*-iKO and Ctrl mice received intraperitoneal tamoxifen injections with 1 mg tamoxifen dissolved in 100 µl corn oil (Sigma Aldrich) with one injection per day on 5 consecutive days, followed by a break of 2 days, and then additional 5 consecutive days of tamoxifen injection.

Genotypes were determined by genomic PCR. Primers to determine the *Ap2m^flox^* allele were sense 5’-CTCATATACG AGCTGCTGGATG, sense 5’-CCAAGGGACC TACAGGACTTC, and antisense 5’-GCTCAAGCCA ACTAGAGTAATC (**Supplemental data Fig. S1a**), amplifying a 265 bp product for the wild type allele, a 441 bp product for the *Ap2m^flox^* allele, and a 372 bp product for recombined *Ap2m^flox^* allele (**Supplemental data Fig. S1b, S1c**), as previously established (Kononenko et al., 2014). Primers to detect the *Cnp^Cre^* allele were sense 5’-GCCTTCAAAC TGTCCATCTC and 5’-CAGGGTGTTA TAAGCAATCCC, and antisense 5’-CCCAGCCCTT TTATTACCAC and 5’-CCTGGAAAAT GCTTCTGTCCG, amplifying a 700 bp product for the WT allele and a 357 bp product for the Cre-allele (Buscham et al., 2022) (**Supplemental data Fig. S1b**). Primers to detect *Plp^CreERT2^* were 5’-TGGACAGCTG GGACAAAGTAAGC and 5’-CGTTGCATCG ACCGGTAATGCAGGC, amplifying a Cre-positive 250 bp product (Leone et al., 2003) (**Supplemental data Fig. S1c**). Mice were bred and kept in the animal facility of the MPI-NAT with 2-5 mice per cage. All mice were housed under a 12 h dark/light cycle with food and water *ad libitum*. All animal experiments were performed in accordance with the German animal protection law and approved by the Niedersächsisches Landesamt für Verbraucherschutz und Lebensmittelsicherheit (LAVES) under license numbers 14/1677, 15/1833, and 19/3334. The animal facility at the MPI-NAT is registered at the LAVES according to TierSchG §11 Abs. 1. According to the German Animal Welfare Law (Tierschutzgesetz der Bundesrepublik Deutschland, TierSchG) and the regulation about animals used in experiments, dated 11^th^ August 2021 (Tierschutz-Versuchstierverordnung, TierSchVersV), an animal welfare officer and an animal welfare committee are established for the institute.

### Biochemical purification of myelin fractions from mouse brains

Myelin-enriched membrane fractions were biochemically purified from mouse brains using sucrose gradient centrifugation and osmotic shocks as described (Erwig, et al. 2019). Brain hemispheres were homogenized in 0.32 M sucrose containing protease inhibitor (Roche Complete, Roche Diagnostics GmbH, Mannheim, Germany) using a T10 ULTRA-TURRAX® homogenizer (IKA, Staufen, Germany). Before myelin purification, a fraction of homogenized brain lysate was taken, snap-frozen on dry ice, and stored at -80°C. Protein concentrations of lysate and myelin samples were measured with the DC Protein Assay Kit (Bio-Rad, Hercules, USA) according to manufacturer’s microplate assay protocol, based on the Lowry assay (Lowry et al., 1951), and optical density was measured at 650 nm using the Eon™ High Performance Microplate Spectrophotometer (BioTek, Vermont, USA) using Gen5 software (BioTek Instruments, Bad Friedrichshall, Germany).

### Myelin proteome analysis

In-solution digestion of myelin proteins by filter-aided sample preparation (FASP) (Erwig et al., 2019a) and LC-MS-analysis by different MS^E^-type data-independent acquisition (DIA) mass spectrometry approaches was performed as recently established for mouse PNS and CNS myelin (Jahn et al., 2020; Siems et al., 2020). Briefly, protein fractions corresponding to 10 μg myelin protein were dissolved in lysis buffer (1% ASB-14, 7 M urea, 2 M thiourea, 10 mM DTT, 0.1 M Tris pH 8.5) and processed according to a CHAPS-based FASP protocol in centrifugal filter units (30 kDa MWCO, Merck Millipore). After removal of the detergents, protein alkylation with iodoacetamide, and buffer exchange to digestion buffer (50 mM ammonium bicarbonate (ABC), 10 % acetonitrile), proteins were digested overnight at 37°C with 400 ng trypsin. Tryptic peptides were recovered by centrifugation and extracted with 40 µl of 50 mM ABC and 40 µl of 1% trifluoroacetic acid (TFA), respectively. For quantification according to the TOP3 approach (Silva et al., 2006), combined flow-throughs were spiked with 10 fmol/μl of yeast enolase-1 tryptic digest (Waters Corporation) and directly subjected to LC-MS-analysis.

Tryptic peptides were separated by nanoscale reversed-phase UPLC and mass spectrometrically analyzed on a quadrupole time-of-flight mass spectrometer with ion mobility option (Synapt G2-S, Waters Corporation) as recently described in detail (Jahn et al., 2020; Siems et al., 2020). The samples were analyzed in the DRE-UDMS^E^ mode in which a deflection device is cycled between full and reduced ion transmission to provide a compromise between the deep proteome coverage of UDMS^E^ and the high dynamic range of MS^E^ (see (Siems et al., 2020) for details). Continuum LC-MS data were processed using Waters ProteinLynx Global Server (PLGS) and searched against a custom database compiled by adding the sequence information for yeast enolase 1 and porcine trypsin to the UniProtKB/Swiss-Prot mouse proteome (release 2016-07, 16806 entries) and by appending the reversed sequence of each entry to enable the determination of false discovery rate (FDR) set to 1% threshold.

For post-identification analysis including TOP3 quantification of proteins, the freely available software ISOQuant (www.isoquant.net) (Distler et al., 2014) was used as described (Jahn et al., 2020; Siems et al., 2020). Only proteins represented by at least two peptides (minimum length seven amino acids, score ≥ 5.5, identified in at least two runs) were quantified as parts per million (PPM), i.e. the relative amount (w/w) of each protein in respect to the sum over all detected proteins. FDR for both peptides and proteins was set to 1% threshold and at least one unique peptide was required. Proteins identified as contaminants from blood (albumin, hemoglobin) or skin/hair cells (keratins) were removed and potential outlier proteins were revised by inspecting the quality of peptide identification, quantification and distribution between protein isoforms. Filtered protein lists were subjected to statistical analysis with the Bioconductor R packages ‘limma’ and ‘q-value’ to detect significant changes in protein abundance by moderated t-statistics as described (Jahn et al., 2020; Siems et al., 2020). Proteome profiling of control and *Ap2m* cKO myelin was performed with three biological replicates per condition and duplicate digestion as technical replicate, resulting in a total of 12 LC-MS runs to be compared. For data visualization by volcano plots, q-values were plotted against fold-change after -log10 and log2 transformation, respectively.

### Immunoblotting

Immunoblotting was performed essentially as described (Gargareta et al., 2022). Brain lysate and purified myelin samples were diluted in 1 X SDS sample buffer (40% Glycerol, 240 mM, Tris/HCl pH 6.8, 8% SDS, 0.04% Bromphenol blue) and 5% Dithiotheitol (DTT). When detecting MAG, samples were prepared without DTT. Before loading the gel, myelin samples were incubated at 40°C and brain lysate samples at 90°C (10 min). Protein separation was performed by SDS-PAGE (8-15% acrylamide) with the Mini-PROTEAN Handcast system (Bio-Rad, Munich, Germany). Depending on the protein of interest 0.5-15 µg of sample was loaded along with 5 µl of the molecular marker PageRuler^TM^ Plus Prestained Protein Ladder (Thermo Fisher Scientific, St. Leon-Rot, Germany). Proteins were separated at a constant current of 180 V for 1 h using a power supply (Bio-Rad, Hercules, USA). Depending on the molecular weight of the protein of interest, immunoblotting was performed with a Novex® Semi-Dry Blotter (Invitrogen, Karlsruhe, Germany) for 45 min at 20 V or a Mini Trans-Blot cell blotting module (BioRad) for 90 min at 100 V. Proteins were transferred to a polyvinylidene difluoride (PVDF) membrane (GE Healthcare Life Science, Chicago, USA) pre-activated in 99% ethanol. The quality of the transfer and samples was analyzed by total protein staining using 1 x Fast Green solution (0.005 mg/ml Fast Green (Merck KGaA, Darmstadt, Germany), 30% Methanol, 6.7% Acetic acid, 63.3% ddH2O). Fast Green signals were detected with the deepRed excitation Filter (670 nm) using the ChemoStar fluorescent imager (INTAS Science Imaging Instruments GmbH, Göttingen, Germany). After Fast Green staining, membranes were blocked in blocking puffer (5% [w/v] non-fat dry milk (Frema instant skimmed milk powder) diluted in 1 x TBS-T (50 mM Tris/HCl, pH 7.5, 150 mM NaCl, 0.05% Tween-20) for 1 h at RT. After blocking, primary antibodies were diluted in blocking buffer and applied to the membrane overnight at 4°C on a horizontal rotor. Following washing in TBS-T (4 x 5 min), membranes were incubated blocking buffer containing the respective diluted horseradish peroxidase (HRP)-coupled secondary antibodies (1 h, RT, 1:10000). Membranes were again washed in TBS-T (4 x 5 min), and bands were visualized using enhanced chemiluminescent detection (ECL) in a 1:1 ratio (Western LightningR Plus-ECL or SuperSignal™ West Femto Maximum Sensitive Substrate; ThermoFischer Scientific, St. Leon-Rot, Germany). Band signals were scanned using the ChemoStar fluorescent imager (INTAS Science Imaging Instruments GmbH, Göttingen, Germany).

Primary antibodies were specific for AP2µ (Abcam, #ab75995, 1:1000), AP2α (BD Bioscience, #AB 397867, 1:500), ATP1a1 (Abcam, #ab7671, 1:5000), CLTC (BD Biosciences, #610500, 1:200), CNP (Sigma Aldrich, #SAB1405637, 1:1000), CTSD (Abcam, #ab75852, 1:1000), LAMP1 (BD Biosciences, #553792, 1:1000), LAMP2 (Abcam, #ab13524, 1:1000), LC3B (Novus Biologicals, #NB100-2220, 1:1000), MAG (clone 513, Chemicon, #MAB1567, 1:1000), MOG (clone 8-18C5, (Linnington et al., 1984), 1:500), p62 (Abnova, #H00008878-M01, 1:500), PLP (A431, (Jung et al., 1996), 1:500). Secondary HRP-coupled antibodies were HRP-goat anti-mouse IgG (Dianova, #115-03-003) (Dianova, Hamburg, Germany), HRP-goat anti-rat IgG (Dianova, #112-035-167), or HRP-goat anti-rabbit IgG (Dianova, #111-035-003).

### Quantitative real time PCR (qRT-PCR)

mRNA abundance was determined by qRT-PCR using dissected corpora callosa. Frozen corpora callosa were homogenized in Trizol (Life Technologies, ThermoFischer Scientific), and RNA was extracted using the Miniprep kit (Qiagen, Portland, USA). The quality and concentration of the isolated RNA was measured using the Nano Drop 2000 spectrophotometer (Thermo Fisher Scientific). The RNA concentration was adjusted to 300 ng/µl, and cDNA was synthesized using random nonamer primers and SuperScriptIII reverse transcriptase (200 U/μl) (Invitrogen, Karlsruhe, Germany). qRT-PCR was performed using SYBR Green PCR Master Mix (Promega, Fitchburg, USA) by a 45 cycle heating protocol in the LightCycler 480 II (Roche Diagnostics GmbH, Mannheim, Germany). Each cDNA sample was tested in triplicates with the respective primers. The mRNA abundance was quantified using the LightCycler 480 Instrument II software and normalized to the mean of the standards *Hprt* and *Rplp0 or Hprt* and *Rps13*. For each sample ΔΔCT values were calculated, and statistical analysis was performed in GraphPad Prism 10 software (Dotmatics, Bishop’s Stortford, UK).

Primers were specific for *Ap2m* (fwd (forward) 5’-CTTGGGATCC GAGAGTGG, rev (reverse) 5’-GTGATGAAGG TTTTCAGTGC, 5’-CACTCTCCAG AACATCTAGG), *Atg5* (fwd 5’-AAGTCTGTCC TTCCGCAGTC, rev 5’-TGAAGAAAGT TATCTGGGTAGCTCA), *Ctsd* (fwd 5’-CCCTCCATTC ATTGCAAGATAC, rev 5’-TGCTGGACTT GTCACTGTTGT), *Lamp1* (fwd 5’-CCTACGAGAC TGCGAATGGT, rev 5’-CCACAAGAAC TGCCATTTTTC), *Lamp2* (fwd 5’-AAGGTGCAAC CTTTTAATGTGAC, rev 5’-TGTCATCATC CAGCGAACAC), *Mag* (fwd 5’-GAAGCCTCTG GAAGAATGTGA, reverse 5’-CACCGTAGAC AGGGTCTTCAA), *Mog* (fwd 5’-ATGAAGGAGG CTACACCTGC, reverse 5’-CAAGTGCGAT GAGAGTCAGC), *Tfeb* (fwd 5’-CCAGAAGCGA GAGCTCACAGAT, rev 5’-TGTGATTGTC TTCTTCTGCCG), *Rplp0* (fwd 5’-GATGCCCAGG GAAGACAG, rev 5’-ACAATGAAGC ATTTTGGATAATCA), *Rps13* (fwd 5’-CGAAAGCACC TTGAGAGGAA, rev 5’-TTCCAATTAG GTGGGAGCAC), and *Hprt* (fwd 5’-TCCTCCTCAG ACCGCTTTT, rev 5’-CCTGGTTCAT CATCGCTAATC). Primers were obtained from the primer synthesis facility (Department of Molecular Neurobiology, MPI-NAT, Göttingen, Germany).

### Electron microscopy

For transmission electron microscopy (TEM), mice were sacrificed by cervical dislocation, the optic nerve and spinal cord removed and directly fixed in Karlsson-Schultz fixative (4% PFA, 2.5% glutaraldehyde in 0.1 M phosphate buffer (PB) (0.36% NaH_2_PO_4_ x H_2_O, 3.1% Na_2_HPO_4_ x 2H_2_O, 1% NaCl)) overnight at 4°C. The samples were prepared and embedded in Epon (Serva, Heidelberg, Germany) resin as described (Weil et al., 2019). After dehydration and infiltration with Epon, the optic nerves were placed in silicon molds filled up with Epon, which was polymerized for 24h at 60°C. The polymerized Epon blocks were sectioned using a PTPC Powertome Ultramicrotome (RMC, Tuscon Arizona, USA) and a DiATOME diamond knife, ultra 35° 3.0mm (Diatome AG, Nidau, Switzerland). Ultrathin sections of 50 nm were cut and collected on formvar polyvinyl-coated copper mesh grids (Science Services, Munich, Germany). To enhance the contrast, sections were stained with Uranyless (Electron Microscopy Sciences, Hatfield, U.S.A) for 30 min followed by 5 washing steps with ddH_2_0. The sections were imaged with a LEO EM912 Omega (Zeiss, Oberkochen, Germany) at 4000 x magnification or a Zeiss EM900 electron microscope at 7000 x magnification. Per mouse, 15-25 non-overlapping images were taken and analyzed in the open source image processing package Fiji (<u>fiji.sc</u>). To quantify the percentage of myelinated and non-myelinated axons, as well as that of pathological profiles, all profiles completely captured on the images were analyzed. All axons were assigned to the following categories: normal-appearing myelinated axon, non-myelinated axon, myelin outfolding, myelin whorl, lamella splitting, axonal spheroid, or other pathology. For g-ratio analysis, the Feret diameter of the axon and the Feret diameter of the corresponding myelin sheath were measured, and the ratio was calculated. Axons for g-ratio analysis were randomly selected by placing a grid (area per point = 5000 µm^2^) on the images. At least 150 axons per mouse were quantified.

### Focused ion beam-scanning electron microscopy

Samples were prepared, cut, and scanned essentially as described (Buscham et al., 2022). Briefly, samples were trimmed with a 90° trimming knife (Diatome AG, Nidau, Switzerland) and attached to a scanning electron microscopy (SEM)-stub (Pin 12.7 mm × 3.1 mm; Science Services GmbH) with a silver filled epoxy (Epoxy Conductive Adhesive, EPO-TEK EE 129–4; EMS, United States) and polymerized at 60° overnight. The samples were coated with a 10 nm platinum layer (sputter coating machine: EM ACE600, Leica) and placed inside the focused ion beam scanning electron microscope (Crossbeam 540, Zeiss). Specimens were coated with a 400 nm deposition of platinum using 3 nA, followed by cross-sectioning the region of interest with a 15 nA ion current and polished with 7 nA. The dataset was acquired at 1.5 kV (1000 pA) with a voxel size of 5 nm × 5 nm × 25 nm and a milling current of 1.5 nA and Atlas 3D (Atlas 5.1, Fibics, Canada) software was used to collect the 3D data. Post-processing was performed using Fiji (Schindelin et al., 2012). Images were cropped, aligned (using the Plugin ‘Linear Stack Alignment with SIFT‘), and inverted. Additionally, the contrast was enhanced using the Contrast Limited Adaptive Histogram Equalization (CLAHE) plugin. Pathological profiles were analyzed in Fiji. All profiles categorized as myelin outfoldings, myelin whorls, or myelin infoldings, in a volume were counted and normalized to a volume of 10000 µm^3^. Exemplified profiles were picked for segmentation and 3D reconstruction using IMOD (v 4.9.12, University of Colorado, USA, bio3d.colorado.edu/imod).

### Cryo-immuno electron microscopy

Immuno-gold labelling of optic nerve cryosections was performed as described (Werner et al., 2007; Siems et al., 2025). Briefly, optic nerves were dissected from *Ap2m* iKO and respective Ctrl mice 6 mo PTI, or from 8 mo old c57Bl/6N wildtype mice. Optic nerves were fixed with fixative (0.25% glutaraldehyde, 4% formaldehyde in 0.1 M phosphate buffer) overnight at 4°C, stained with toluidine blue (Merck, Darmstadt, Germany) and embedded in 10% gelatin (Twee Torens, Delft, The Netherlands) blocks. Gelatin blocks were infiltrated with sucrose in 0.1M phosphate buffer overnight (4°C), mounted on aluminum pins (Senkniete DIN661) and frozen in liquid nitrogen. Cryo-sections of 50 nm were sectioned using a Diatome Diamond knife, cryo-immuno 2,0 mm (Diatome, Biel, Switzerland) on a Leica UC6 ultramicrotome with FC6 cryochamber (Leica, Vienna, Austria). The sections were transferred to carbon coated grids (Science Services, Munich, Germany) and stored at 4°C. Before antibody labelling, the sections were washed in PBS (1.7 M NaCl, 34 mM KCl, 40 mM Na_2_HPO_4_ x 2H_2_O, 18 mM K_2_HPO_4_, pH 7.2 with 1 N NaOH) (3 x 2 min), followed by two washing steps in PBS with glycine (0,1%), and blocked in 1% BSA in PBS (3 min). Primary antibodies were diluted in 1% BSA in PBS and applied for 1 h. The sections were then washed and incubated with the respective Immuno-gold coupled secondary antibody (1:40). Sections were washed again (5 x 2 min, PBS) and fixed with 1% glutaraldehyde in PBS (5 min). To enhance the contrast, the sections were stained with uranyl oxaliacetate (4%, pH 7) and methylcellulose-uranyl acetate (4% uranyl acetate (Merck, Darmstadt, Germany) diluted 1:10 in 2% methylcellulose (Sigma-Aldrich, Steinheim, Germany), and imaged with a LEO EM912 Omega (Zeiss, Germany).

Primary antibodies were specific for AP2µ (α-AP50, BD Biosciences, #610502, 1:200), CLTC (BD Biosciences, #610500, 1:200), LAMP1 (BD Biosciences, #553792, 1:300), or LAMP2 (Abcam, #ab13524, 1:200). Secondary immuno-gold coupled antibodies were α-rat Immuno-gold (Aurion, #810.455) or α-mouse Immuno-gold (Aurion, #810.422) (Aurion, Wageningen, The Netherlands).

### Immunohistochemistry

For immunohistochemistry, mice were transcardiacally perfused with Hank’s buffered salt solution (HBSS, Gibco, Thermo Fisher Scientific) and 4% paraformaldehyde (PFA) in 0.1 M phosphate buffer for tissue fixation. The tissue was dissected, and post-fixed with 4% PFA in 0.1 M phosphate buffer overnight and stored in PBS until further processing. For fluorescent immunolabelling of spinal cord sections, the perfused spinal cords were transferred into 10 % [w/v] sucrose in 0.1 M PB for 1 h, followed by incubations in 20 % [w/v] sucrose in 0.1 M PB and 30% [w/v] sucrose in 0.1 M PB for one night each. Samples were embedded using Tissue-Tek® O.C.T.TM Compound (Sakura Finetek, Torrance, USA) and stored at -20°C. 10 μm cross section of spinal cords were cut with a CM1950n cryostat (Leica Biosystems, Wetzlar, Germany), collected and transferred to Superfrost® Plus microscope slides (Thermo Fischer Scientific). Sections were first post-fixed for 3 min with ice cold 4% PFA followed by 3 min permeabilization with ice-cold methanol. After washing with PBS (3 x 5 min) the sections were blocked with blocking buffer (10% horse serum, 0.25% Triton X-100, 1% bovine serum albumin in PBS; 1h, RT). Primary antibodies were diluted in 1.5% horse serum, 0.25% Triton X-100 in PBS and applied overnight at 4°C. Primary antibodies were specific for LAMP1 (BD Biosciences, #553792, 1:1000) and MBP ((Meschkat et al., 2022), 1:1000). Samples were washed with PBS (3 x 5 min), and secondary antibodies diluted in PBS were applied together with 4‘,6-diamidino-2-phenylindole (DAPI; 1:50000 in PBS) for 1 h at RT. After another two washing steps in PBS (5 min) the samples were mounted using Aqua-Poly/Mount (Polysciences,). Fluorescent sections were captured using Zeiss Observer Z1/Z2 microscopes equipped with a Colibri 5 LED light source (630 nm, 555 nm, 475 nm, 385 nm excitation) and Zeiss Filter sets: 96 HE BFP, 90 HE DAPI/GFP/Cy3/Cy5, 38 GFP, 43 DsRed, and 50 Cy5. Imaging was performed with an Axiocam MrM (Zeiss) at magnification of 20x, 40x, or 63x. Additionally, high resolution imaging was performed using a Leica SP8 Lightning confocal microscope (Leica Biosystems), featuring an adjustable white-light laser, at 63 x magnification. To display the images, channels were assigned pseudo-colors and adjusted using Fiji.

Chromogen labeling to analyze neuropathology and oligodendrocyte numbers was performed as described (De Monasterio-Schrader et al., 2013; Lüders et al., 2017). Briefly, fixed brain samples were embedded in paraffin using an automated embedding system (Leica) and sectioned into 5 µm sections using the Microm HM400R Microtome (Thermo Fisher Scientific Waltham, USA). Primary antibodies were specific for the oligodendrocyte marker aspartoacylase (ASPA) (US Biological, #MBS422542, 1:4000), the astrocyte marker glial fibrillary acidic protein (GFAP) (Novo Castra, #NCL-L-GFAP-GA5, 1:200), the microglia marker ionized calcium-binding adapter molecule 1 (IBA1) (abcam, #ab5076, 1:1000), or amyloid beta precursor protein (APP) (Chemicon, #MAB348, 1:1000) as a marker for axonal swellings. IBA1 and ASPA antibodies were detected using the Vector Elite ABC kit (Vector Labs, Newark, Germany); GFAP and APP antibodies were detected using the LSAB2 kit (Dako, Glostrup, Denmark). The slides were imaged using an AxioImager Z1 bright-field light microscope (Zeiss, Oberkochen, Germany) coupled to an AxioCam MRc Camera (Zeiss, Oberkochen, Germany) and controlled by Zeiss Zen 1.0 software (Zeiss, Oberkochen, Germany). Images were taken at 40 x magnification, stitched using Zeiss Zen 1.0 software, and further processed using Fiji. ASPA-immunopositive cells and APP-immunopositive axonal swellings were counted manually and normalized to the quantified area. APP-immunopositive labelling was only quantified as axonal swelling if the minimum size was at least half of the size of a nucleus (marked with DAPI) and not in immediate proximity with a cell body. To quantify IBA1- or GFAP immunopositive area, an ImageJ plug-in was used based on color threshold and immuno-positive area was normalized to the quantified area as reported (De Monasterio-Schrader et al., 2013; Lüders et al., 2017).

### Preparation of primary oligodendrocyte cultures

Primary oligodendrocytes cultures were prepared from wild type c57Bl/6N mouse brains at postnatal day 8 (P8) as reported (Weil et al., 2019; Sasmita et al., 2024; Späte et al., 2024). In brief, oligodendrocyte precursor cells were isolated using magnetic activated cell sorting and anti-NG2 MicroBeads (Miltenyi Biotec, Bergisch Gladbach, Germany). For brain dissociation, the neural tissue dissociation kit was used according to the manufacturer’s protocol. Brains were dissected, cut in pieces and transferred into tubes containing enzyme mix 1 and incubated (37°C for 15 min). After the first incubation, enzyme mix 2 was added followed by another incubation step (37°C, 20 min) with rotation in between. Following tissue dissociation, the tubes were centrifuged (5 min, 1200 rpm), the supernatant discarded, and the pellet resuspended in DMEM (Gibco, Thermo Fisher Scientific) with 1% horse serum. The cell suspension was passed through a 70 μm cell strainer followed by another separation through a 40 µm strainer. Tubes were again centrifuged (1200 rpm, 10 min), and the pellet was resuspended and incubated in prewarmed OPC culture medium (100 ml NeuroMACS media, 2 ml MACS NeuroBrew21, 1 ml Pen/Strep, 1 ml L-GlutaMAX; 37°C, 2 h). Following the incubation in OPC culture medium, tubes were centrifuged (1200 rpm, 10 min, 4°C), and the pellet incubated in NG2 MicroBeads diluted in DMEM (10 µl NG2-beads per 10^7^ total cells, 15 min, 4°C). The cell suspension was again centrifuged (1200 rpm, 10 min, 4°C), the supernatant removed, and the pellet resuspended in 5 ml DMEM with 1% horse serum. LS columns (Miltenyi Biotec) were attached to a magnet and activated with DMEM. After adding the cell suspension to the LS columns, the columns were washed 3 x with DMEM. To collect the NG2 bound cells the columns were detached from the magnet and flushed with 5 ml DMEM. OPC were plated at a density of 12 x 10^4^ cells per 12 well plate, cultured in proliferation medium, which after 4 days *in vitro* (DIV 4) was replaced with OPC differentiation medium. Cells were fixed at DIV 7 with 4% PFA, and washed with PBS (3 x 5 min). For fluorescent immunolabelling, cells were permeabilized with ice-cold 2% Triton X-100, and blocked with blocking buffer (10% horse serum, 0.25% Triton X-100, 1% bovine serum albumin in PBS; 1h, RT). Primary antibodies specific for AP2α (AP-6, Invitrogen #MA1-064, 1:200) or MBP (Meschkat et al., 2022, 1:1000) were diluted in incubation buffer (1.5% horse serum in PBS) and applied for 4 h at 4 °C. Coverslips were washed with PBS (3 x 5 min) and incubated in secondary antibodies (1 h) diluted in PBS (α-rabbit STAR-RED, abberior, 1:200; α-mouse STAR-ORANGE, abberior, 1:200). After washing in PBS (2 x 5 min) the samples were incubated with 4‘,6-diamidino-2-phenylindole (DAPI; 1:50000 in PBS). Cells were briefly washed again with PBS and mounted using abberior mount solid mounting medium (Abberior Instruments, Göttingen, Germany). Images were obtained using a Confocal and STED FACILITY line microscope (Abberior Instruments) equipped with a 60x oil immersion objective (N.A. 1.42, Olympus, Japan).

### Elevated Beam assay

The elevated beam test (Luong et al., 2011; Berghoff et al., 2021) was used to determine motor capabilities of *Ap2m* iKO and littermate control mice 9-10 mo PTI. Briefly, mice cross a 55 cm long and 1 cm wide beam with a slight slope and was located 50-60 cm above the tabletop (custom made, MPI of Multidisciplinary Sciences). A shelter box was placed at the far end of the beam to serve as motivation. Mice were placed in their home cage after reaching the box. The time a mouse needed to cross the beam and the number of slips were measured. Mice were habituated for 2 training days and tested on the following third day. On each day the mice had to cross the beam for 3 times. After each mouse the device was cleaned with ethanol and water. The average number of slips and crossing time per mouse was calculated from all three runs on the test day. One control mouse was identified as an outlier via the GraphPad Prism online tool (www.graphpad.com/quickcalcs/Grubbs1.cfm) and excluded from the analysis.

### Quantifications and statistical analysis

All experiments were analyzed blinded for the genotypes. Statistical assessment was performed with GraphPad Prism 10 or R-studio. To compare two groups, unpaired students t-test was applied with Welch‘s correction if samples were unequally distributed. To compare more than two groups, two-way ANOVA was applied as indicated in the figure legends. Significance levels are represented as ns = not significant, *p<0.05, **p<0.01, ***p<0.001 with exact p-values indicated in the figure legends. Mouse numbers are indicated in the figures or figure legends. Outlier tests were performed using the GraphPad Prism online tool (www.graphpad.com/quickcalcs/Grubbs1.cfm); outliers were only excluded where indicated. Calculations and illustration of data was performed using Fiji, Microsoft Excel 2013, GraphPad Prism 10 (GraphPad Software Inc, San Diego, USA), and Adobe® Illustrator 2022 (San Jose, USA).

## DATA AVAILABILITY STATEMENT

All relevant data are included in the main paper or supplemental files. The mass spectrometry proteomics data have been deposited to the ProteomeXchange Consortium via the PRIDE (Perez-Riverol et al., 2025) partner repository and will be accessible with dataset identifier PXD060968 upon publication in a journal.

## ACKNOWLEDGMENTS

We thank K. Bergann, A. Fahrenholz, K. Gitt, D. Hesse, and T. Ruhwedel for technical support, P. Saftig and J. Büscher for discussions, L. Piepkorn for help with statistics, K. Kusch and C. Linington for antibodies, and the International Max Planck Research School for Genome Science (IMPRS-GS) for supporting S.B.S. and S.H.. K.-A.N. holds an European Research Council (ERC) Advanced Grant (‘MyeliNano‘ to K.-A.N.). This work was supported by the Deutsche Forschungsgemeinschaft (DFG grant WE 2720/2-2 to H.B.W.). Open access publication funding and organized by project DEAL.

## AUTHOR CONTRIBUTIONS

SBS Investigation, Data analysis, Writing-original draft, Writing-review & editing

RBJ Investigation, Writing-review & editing

OJ Methodology, Data analysis, Writing-review & editing

MM Investigation, Writing-review & editing

SM Investigation, Writing-review & editing

NL Investigation, Writing-review & editing

SH Investigation, Writing-review & editing

AOS Investigation, Writing-review & editing

WM Methodology, Data analysis, Supervision, Writing-review & editing

FB Methodology, Writing-review & editing

NB Supervision, Writing-review & editing EMKA Supervision, Writing-review & editing

VH Conceptualization, Writing-review & editing

KAN Conceptualization, Writing-review & editing

HBW Conceptualization, Funding, Supervision, Writing-original draft, Writing-review & editing

## Figures and Supplementary Material

**Supplementary Table 1:** Myelin proteome analysis dataset

**Supplementary Video 1:** Three-dimensional reconstruction of representative myelin outfoldings and whorls in a tissue block of optic nerve dissected from an *Ap2m* iKO mouse 6 mo PTI and assessed by FIB-SEM, as in Fig. 3i**-3m**. Shown is one mouse representative of three *Ap2m* iKO mice. Myelin is false-colored in green; the corresponding axon is false colored in blue.

**Supplementary Video 2:** Three-dimensional reconstruction of representative normal-appearing axon/myelin-units in a tissue block of optic nerve dissected from a Ctrl mouse 6 mo PTI and assessed by FIB-SEM, as in Fig. 3i**-3m**. Shown is one mouse representative of three Ctrl mice. Myelin is false-colored in green; the corresponding axon is false colored in blue.

---

**Supplementary Figure S1:**
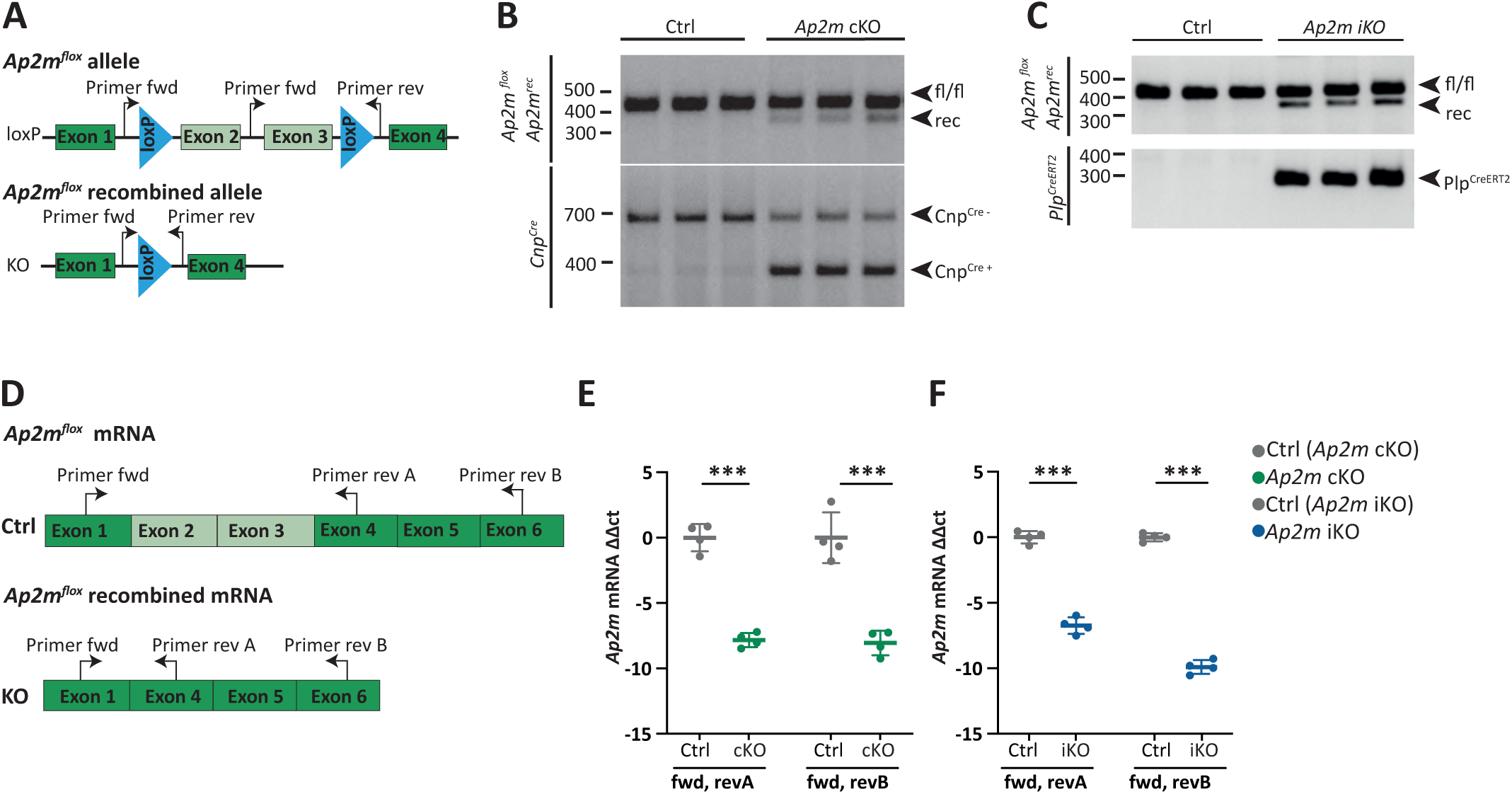
Recombination of *Ap2m* in oligodendrocytes. **A** Scheme of the engineered *Ap2m^flox^* allele (Kononenko et al., 2014), in which exon 2 and 3 are flanked by loxP-sites to allow *Cre*-mediated recombination. Arrows mark positions of genotyping PCR primers. Fwd, forward, rev, reverse. **B,C** Routine genotyping PCR using genomic DNA isolated from optic nerves of *Ap2m^flox/flox^;Cnp^Cre/WT^* (cKO) (**B**), *Ap2m^flox/flox^;Plp^CreERT2^* (iKO) (**C**), and respective Ctrl mice. **B** The upper gel shows the PCR product detecting the *Ap2m^flox^* and the recombined (rec) allele. The lower gel shows PCR products indicating the *Cnp* allele (*Cnp^Cre^*, *Cnp^WT^*). n=3 per genotype. **C** The upper gel shows the PCR product detecting the *Ap2m^flox^* and the recombined (rec) allele. The lower gel shows the PCR product detecting *Plp^CreERT2^*. *Ap2m^flox^, Cnp^Cre^*, and *Plp^CreERT2^* mice were previously published (Kononenko et al., 2014; Leone et al., 2003; Lappe-Siefke et al., 2003). **D-F** qRT-PCR to assess *Ap2m* transcript abundance. **D** Scheme of *Ap2m* transcript with or without *Ap2m^flox^* recombination. Arrows mark positions of qRT-PCR primers. **E,F** mRNA was isolated from corpus callosum of *Ap2m*-cKO and Ctrl mice at P17 (**E**) and of *Ap2m*-iKO and Ctrl mice 3 mo PTI (**F**) and subjected to qRT-PCR. Note that *Ap2m* mRNA displays strongly reduced ΔΔct values in *Ap2m*-cKO and *Ap2m*-iKO compared to Ctrl mice. Data show mean ±SD; datapoints indicate individual mice; n=4; two-tailed unpaired t-test: **E** fwd & rev A p<0.0001, fwd & rev B p=0.0003, F fwd & rev A p<0.0001, fwd & rev B p<0.0001.

**Supplementary Figure S2:**
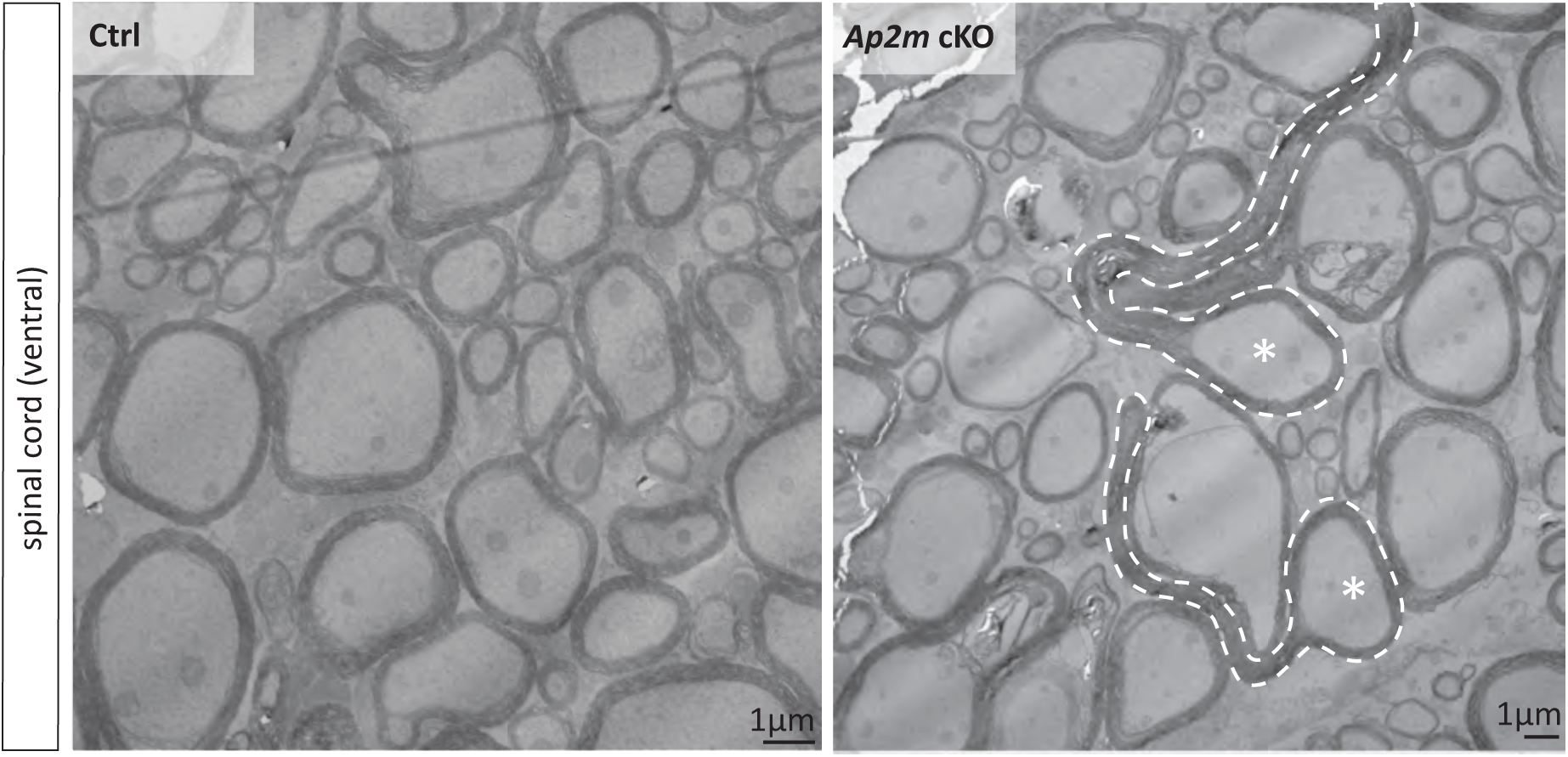
Myelin outfoldings in the spinal cord of *Ap2m-*cKO mice. Representative electron micrographs of spinal cord cross sections from Ctrl and *Ap2m*-cKO mice, as exemplified for P24. Myelin outfoldings are marked with stippled lines; the asterisks mark associated axons. Note that myelin outfoldings were an evident neuropathological feature in *Ap2m*-cKO spinal cords (not quantified). Shown is one mouse per genotype representative of n=4-5 mice per genotype. Scale bar, 1µm. For optic nerves see Fig. 2c**-2i**.

**Supplementary Figure S3:**
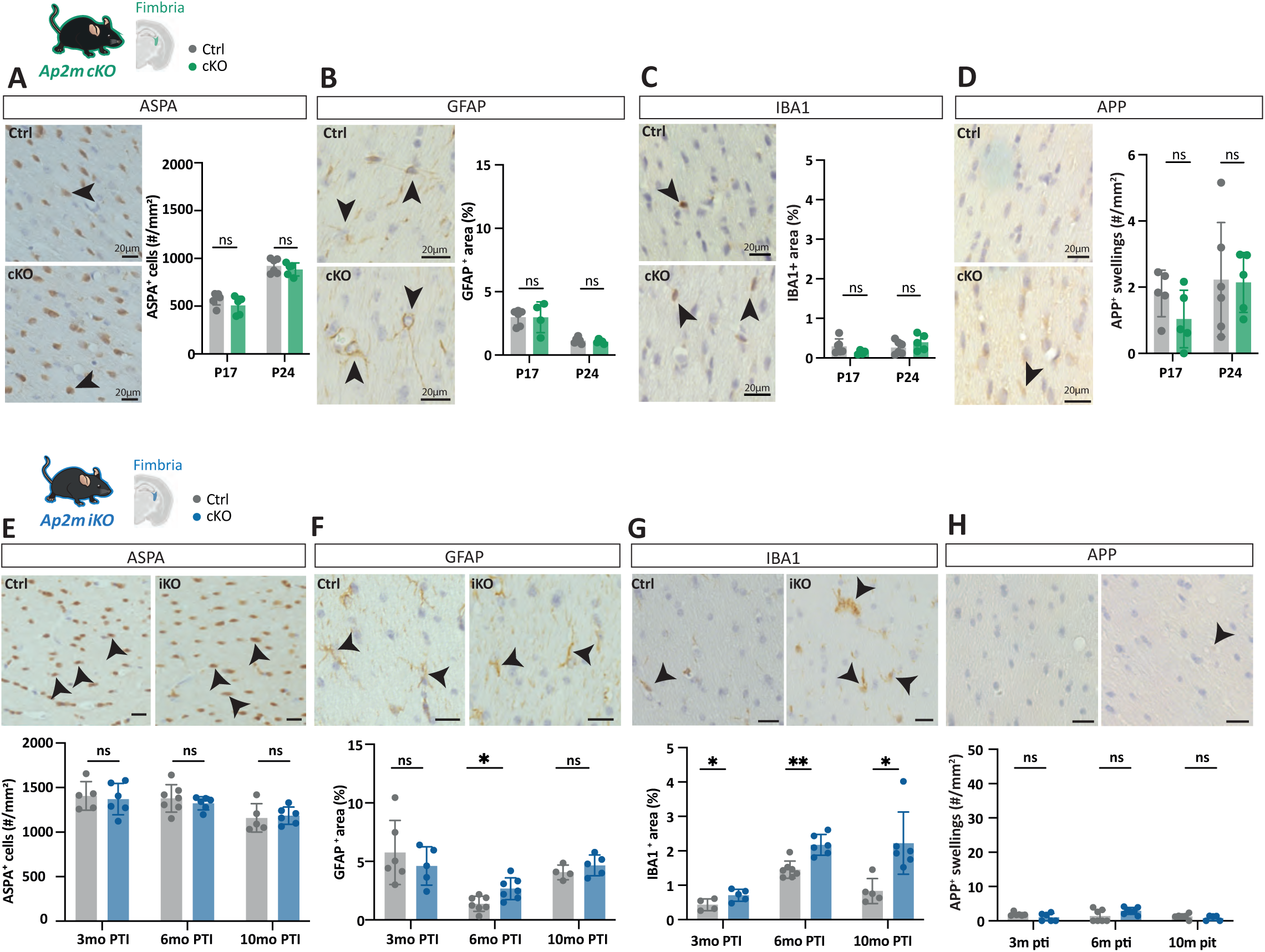
Neuropathological analysis of white matter in *Ap2m*-cKO and *Ap2m*-iKO mice. **A-H** Representative light micrographs of a white matter tract (hippocampal fimbria) of *Ap2m*-cKO mice at P24 (**A-D**) and of *Ap2m*-iKO mice 6 mo PTI (**E-H**) immunolabelled for the oligodendrocyte cell body marker ASPA (**A,E**), the astrocyte marker GFAP (**B,F**), the microglia marker IBA1 (**C,G**), and the marker amyloid precursor protein (APP) to label axonal swellings (**D,H**). Black arrowheads point at immunopositive cells. Scale bars 20 µm. **A-D** Genotype-dependent quantification of the number of ASPA-immunopositive cell bodies (**A**), GFAP-immunopositive area (**B**), IBA1-immunopositive area (**C**), and number of APP-positive swellings (**D**) in *Ap2m* cKO and Ctrl mice at P17 and P24. Data shown as mean ±SD. Datapoints indicate individual mice; n=4-6 per genotype and age. Unpaired t-test with Welch’s correction **A** P17 p=0.207, P24 p=0.430; **B** P17 p=0.988, P24 p=0.480; **C** P17 p=0.992, P24 p=0.286; **D** P17 p=0.162, P24 p=0.922.**E-H** Genotype-dependent quantification of the number of ASPA-immunopositive cell bodies (**E**), GFAP-immunopositive area (**F**), IBA1-immunopositive area (**G**), and number of APP-positive swellings (**H**) in *Ap2m*-iKO and Ctrl mice 3 mo, 6 mo and 10 mo PTI. Note that IBA1-immunopositive area was moderately increased in *Ap2m*-iKO compared to Ctrl mice at all timepoints. Data shown as mean ±SD. Datapoints indicate individual mice; n=5-7 per genotype and age. Unpaired t-test with Welch’s correction **E** 3 mo PTI p=0.732, 6 mo PTI p=0.423, 10 mo PTI p=0.400; **F** 3 mo PTI p=0.415, 6 mo PTI p=0.012, 10 mo PTI p=0.283; **G** 3 mo PTI p=0.049, 6 mo PTI p=0.001, 10 mo PTI p=0.029; **H** 3 mo PTI p=146, 6 mo PTI p= 0.08, 10 mo PTI p= 0.367.

**Supplementary Figure S4:**
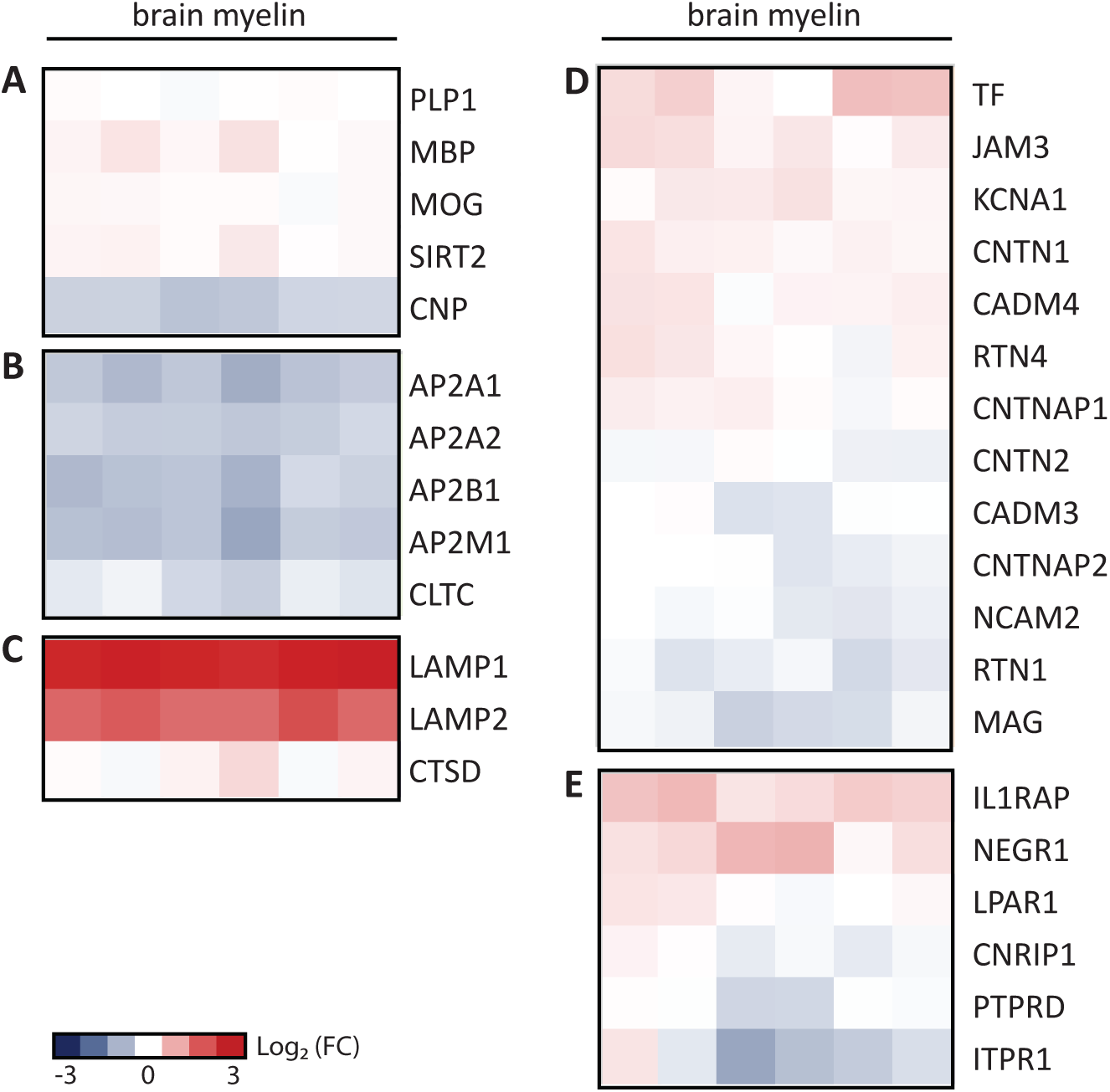
Heatmap comparing the abundance of proteins in *Ap2m*-cKO and Ctrl myelin. Differential quantitative proteome analysis of myelin purified from the brains of *Ap2m*-cKO and Ctrl mice at age P24. Heatmaps show mass spectrometric quantification of selected proteins as measured in three biological replicates of *Ap2m*-cKO myelin with two technical replicates each plotted on a log2-color scale; Each horizontal line displays the fold-change (FC) of a protein in *Ap2m*-cKO myelin compared to its mean abundance in Ctrl. Red represents higher abundance in *Ap2m*-cKO myelin, blue represents higher abundance in control myelin. Shown are selected known myelin proteins **(A)**, constituents of the AP2 complex and clathrin heavy chain (CLTC) **(B)**, lysosomal proteins **(C)**, proteins of the axon/myelin interface **(D)**, and receptors **(E)** including associated proteins present in the dataset. Note the comparatively strong increase in the abundance of lysosomal-associated membrane proteins (LAMP1, LAMP2) in *Ap2m*-cKO myelin. For volcano plot visualization of entire dataset see Figure 4a; for dataset see **Supplementary Table 1**.

**Supplementary Figure S5:**
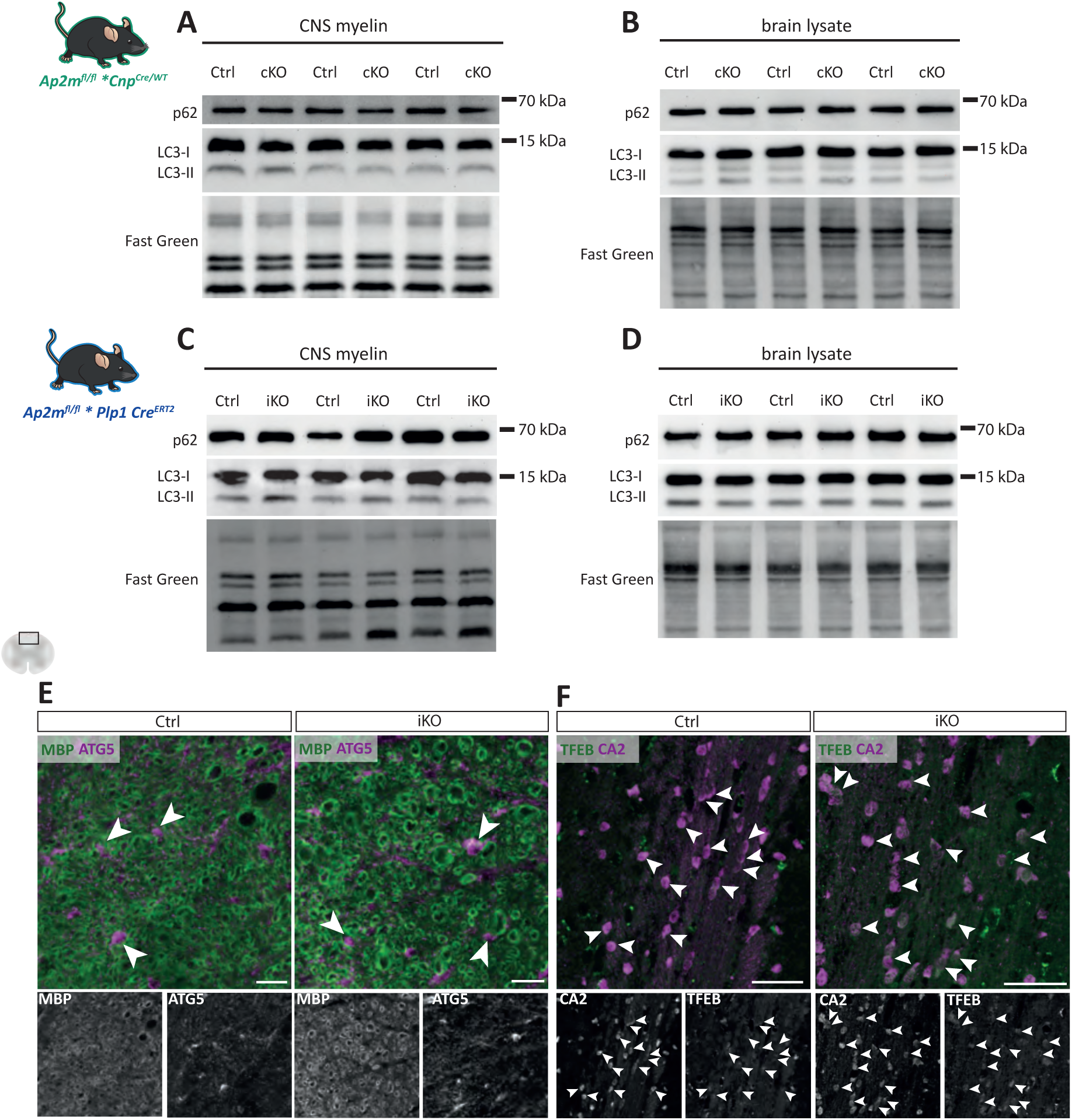
Absence of evidence for induction of autophagy in *Ap2m*-cKO and *Ap2m*-iKO mice. **A-D** Immunoblot analysis of the autophagosomal proteins p62 and LC3 in purified myelin (**A,C**) and brain lysates (**B,D**) of *Ap2m*-cKO (**A,B**), *Ap2m*-iKO (**C,D**) and respective Ctrl mice does not indicate genotype-dependent differences. Fast green protein staining serves as loading and transfer control. Blots show n=3 per genotype; **A,B** P24, **C,D** 6 mo PTI. **E** Immunohistochemical analysis of spinal cord cross sections from *Ap2m-*iKO and Ctrl mice immunolabelled for the autophagy marker ATG5 (magenta) and the myelin marker MBP (green). Note that there is no abundance change in ATG5 immunopositive structures associated with myelin profiles. Scale bar, 20µm. **F** Immunohistochemical analysis of spinal cord cross sections from *Ap2m-*iKO and Ctrl mice immunolabelled for the autophagy-inducing transcription factor TFEB (magenta) and the oligodendrocyte marker CA2 (green). Scale bar 50 µm.

